# A Differentially Methylated CpG Site in the *IL4* Gene Associated with Anti-FVIII Inhibitor Antibody Development in Hemophilia A

**DOI:** 10.1101/550566

**Authors:** Thiago Barbosa de Souza, Thais Louvain de Souza, Cristina dos Santos Ferreira, Cleiton Figueiredo Osório da Silva, Liliana Carmen Rossetti, Vanina Daniela Marchione, Carlos Daniel De Brasi, Enrique Medina-Acosta

**Affiliations:** Laboratório de Biotecnologia, Núcleo de Diagnóstico e Investigação Molecular, Centro de Biociências e Biotecnologia, Universidade Estadual do Norte Fluminense Darcy Ribeiro, Brazil; Faculdade de Medicina de Campos, Campos, Brazil; Instituto de Medicina Experimental, CONICET, Academia Nacional de Medicina de Buenos Aires, Argentina; Instituto de Investigaciones Hematológicas Mariano R Castex, Academia Nacional de Medicina, Argentina

**Keywords:** anti-FVIII inhibitor antibody, DNA methylation, 5meCpG, *F8* Inv22 inversion, hemophilia A, rs35081782, rs2227282

## Abstract

Hemophilia A is the most common clotting disorder in humans. It affects one in five thousand live-born children. Mutations in the X-chromosome linked *F8* gene lead to the deficiency of circulating factor VIII (FVIII). The defect is characterized by severe bleeding. The standard therapy is to replace the deficient factor intravenously. The main adverse event of the therapy is the development of anti-FVIII inhibitor antibodies that impair coagulation and result in increased complications and risk of death. Several risk factors have been described for the development of inhibitor antibodies, among them age, type of FVIII administered, ethnicity, and variant alleles in immune response genes. Epigenetic risks factors have not yet been explored. This work aimed to evaluate the methylation statuses at thirteen CpG sites (5meCpG) in regulatory regions of the *IL1B, IL2, IL4, IL6, IL10, TNF, IFNG, CTLA4, CD28*, and *CST7* immune regulation genes in hemophilia A affected males on replacement therapy who develop or do not develop inhibitor antibodies. At each of the thirteen specific CpG sites, we observed one of three possible statuses: hypermethylated, hypomethylated or intermediate methylated. We found a statistically significant (*p* = 0.04) decrease in the methylation level at one CpG site in the *IL4* intron 1 (CpG-3) in the affected group of patients presenting with anti-FVIII inhibitors as compared with the group of patients without inhibitors. The differential 5meCpG-3 maps within a predicted enhancer region in *IL4* intron 1 that overlaps DNase I hypersensitive chromatin region of the Th_2_ *IL5, IL13*, and *IL4* cytokine gene cluster and, therefore, permissive for gene expression. Six-bp upstream of the differentially 5meCpG-3 is the rs2227282 *cis* expression quantitative trait locus that influences the transcript levels of the *PDLIM4, SLC22A4, SLC22A5, RAD50, IL4, KIF3A, SEPT8* genes. We consider the *IL4* (CpG-3) site a promising lead epigenetic mark, the potential value of which must be appraised in a larger group of patients. The methodology employed also allowed to evaluate the distribution of the *IL6* rs35081782 insertion/deletion variant, associated with white blood cell count traits in genome-wide association studies, and which showed no difference in distribution between the groups of patients.

## Introduction

Hemophilia A (MIM 306700) is the most severe and frequent blood clotting disorder in humans (Graw et al., 2005). Mutations in the *F8* gene, responsible for the production of clotting factor VIII (FVIII), cause the disease. The FVIII deficiency gives rise to bleeding episodes of challenged control during trauma or spontaneously. One of the most frequent mutations that causes hemophilia A is the inversion of the *F8* intron 22 (Inv22) (Ghosh and Shetty, 2009). This mutation occurs in about 45% of cases (Antonarakis et al., 1995; Rossetti et al., 2011). Among the other mutations are deletions, insertions and point mutations that result in stop-codons or amino acid exchanges (Margaglione et al., 2008; Oldenburg et al., 2010; Rallapalli et al., 2013; Rydz et al., 2013). As standard therapy, hemophiliacs are treated with intravenous infusion of the deficient factor, in recombinant or plasma-derived form (Carcao and Lambert, 2010). The main adverse event during replacement therapy is the development of anti-FVIII inhibitory antibodies (Witmer and Young, 2013; Santagostino et al., 2018). Inhibitor development relates to FVIII antigen presentation and polyclonal activation of T lymphocytes (Astermark, 2006; Witmer and Young, 2013). Inhibitors are polyclonal alloantibodies of the IgG type (IgG1 and IgG4) that neutralize either endogenous or exogenous FVIII activity (Montalvao et al., 2015). Patients who develop inhibitors have a Th_2_-polarized immune response, whereas individuals who do not develop inhibitors exhibit a predominantly Th_1_ response (Chaves et al., 2010a; Chaves et al., 2010b; Oliveira et al., 2013).

Risk factors for inhibitor development are classified into non-genetic and genetic. Among non-genetic risks, the age at the replacement therapy, the type of hemorrhage and the factor administered have been described (Ragni et al., 2009). Among the genetic risk factors, the most well-described factor is the type of causative mutation. Mutations that associate with a higher risk of developing inhibitors are linked to significant functional alterations or complete absence of FVIII. About 40% of patients with large deletions develop inhibitors, while nonsense mutations account for up to 30% of patients (Zhang et al., 2009). Several studies have sought to establish a relationship between genetic variants of the immune response and inhibitors, but a few reported statistical significance (Astermark et al., 2006a; Astermark et al., 2006b; Astermark et al., 2007; Bafunno et al., 2010). An association between the promoter SNP rs5742909 (NC_000002.11:g.204732347C>T) in position −318bp in the *CTLA4* (Cytotoxic T-lymphocyte associated protein 4) gene, a crucial T-cell regulator, was also found, where the T allele associated with protection against the development of inhibitors (Astermark et al., 2007; Pavlova et al., 2008). A study in Brazilian patients evaluated the influence of haplotypes across three SNPs in the *IL10* gene promoter region: rs1800896 (NC_000001.10:g.206946897T>C, at −1082bp), rs1800871 (NC_000001.10:g.206946634A>G, at −819bp), and rs1800872 (NC_000001.10:g.206946407T>G, at −592bp). The study found that individuals bearing the CGG or TGG haplotypes are 5.82 times more likely to develop inhibitors, whereas the CGG or TAT haplotypes appear to confer protection because they are more frequent in the group of patients without inhibitors (Chaves et al., 2010a). A genome-wide association study (GWAS) showed a protective role for the missense variant rs3754689 (C_000002.11:g.136590746C>T) in the lactase *LCT* gene (Gorski et al., 2016). The role of the *LCT* gene (i.e., hypolactasia) in the immune response remains unclear. Another study (Astermark et al., 2013) associated thirteen SNPs with p < 0.001, eight of these showed protective effects compared with five exhibiting risks for inhibitor development. The five SNPs associated with inhibitor development are on genes *CD44, CSF1R, DOCK2, MAPK9* and *IQGAP2*.

An association between a polymorphic microsatellite locus in the promoter region of the *IL10* gene, an essential regulatory cytokine, has also been reported (Astermark et al., 2006b). This variant consists of a repeat of the two nucleotides [CA]_n_. The allele with [CA]_n = 19_was associated with inhibitors in severe hemophiliac A due to the Inv22 mutation. Moreover, a comparative gene expression study using microarray analysis identified 545 differentially expressed genes in hemophilia A patients with an inhibitor as compared to non-inhibitor patients, with notable attention for the *IL8* gene (Hwang et al., 2012). Another study identified the small ncRNA mir-1246 with six-fold higher expression in hemophilic patients without inhibitors (Sarachana et al., 2015). As seen, several risk factors have been described for the development of inhibitors, but few studies are conclusive about the causation or impact of genetic variability on this phenomenon.

Surprisingly, no reports are assessing the methylation statuses of the promoter regions or around the SNPs or genes that so far have been associated with the risk of developing or the protection against the production of inhibitor antibodies. The (epi)genetic hallmarks underlying the functional phenotypic difference in immune response between hemophilia A patients who produce and those who not produce anti-FVIII inhibitor antibodies remains, therefore, primarily uncovered. The purpose of this work was to evaluate in hemophiliac patients with and without inhibitors the state of methylation of individual CpG sites located in genes known to modulate the antibody immune response to increase our understanding of the epigenetic mechanisms involved in the inhibitor development. We reasoned that epigenetic marks in the form of differentially methylated 5meCpG sites at the immune genes known to participate in the regulation of antibody production might account in part for the phenotypic variation observed in patients with hemophilia A who develop anti-FVIII antibody inhibitors as compared with those who do not produce them.

## Materials and Methods

### Ethical Aspects

The Ethics Committee of the Institutes of the National Academy of Medicine of Buenos Aires, Argentina (CEIANM) approved the study (June 12, 2013). Peripheral blood samples from participating healthy individuals (n = 20), and hemophilia A subjects (n = 40, being n = 20 with anti-FVIII antibody inhibitors and n = 20 without inhibitors) were collected with written informed consent. The genomic DNA bank was from a single hemophilia center in Argentina. Because of the scarcity and value of the DNA samples from the hemophilia A subjects, for the standardization of the methylation assays, we tested blood DNA samples from healthy Brazilian subjects (n = 10) from a related research project approved by the Ethics Committee of the Faculdade de Medicina de Campos (May 13, 2010). All the participants or legal guardians gave written informed consent.

### Study Design

A case-control study (with or without inhibitor) involving patients with severe hemophilia A. The genomic DNA bank repository consisted of samples from hemophilia A patients stratified about the type of causative mutation. The participants were 40 men with severe hemophilia A caused by Inv22, classified by inhibitor status in subjects positive for inhibitors (n = 20), including index cases with high response (+HR, > 5 UB/mL), and low response (+LR, 5 UB/mL), and individuals who were negative (-) for inhibitors (n = 20), including negative cases (< 0.5UB/mL), and transients whose inhibitor titers disappear before a six-month period (Supplemental Table S1). The patients had been genotyped earlier as positive for either the Inv22-1 or Inv22-2 causative mutations by inverse shifting-PCR (Rossetti et al., 2005; Radic et al., 2009; Rossetti et al., 2011). We also included a group of healthy males (n = 20) from the same ethnic background than the patients from the case and control groups (Supplemental Table S2).

### DNA Extraction

Genomic DNA was extracted from peripheral blood samples using the DNA extraction, RNA and Allprep DNA RNA Protein Mini Kit (QIAGEN) extraction kit according to the manufacturer’s specifications.

### Determination of Methylation Point Profiles

The methylation level at the individual CpG sites was estimated using methylation-sensitive restriction enzyme PCR (MSRE-PCR) triplex specific assays and the products analyzed by Quantitative Fluorescent PCR. We have outlined the concept of the MSRE-PCR triplex assay for the identification of differentially methylated regions in the context of genomic imprinting (Alves da Silva et al., 2016; de Sa Machado Araujo et al., 2018). Briefly, for the intended individual CpG sites herein, each CpG sites was interrogated by a specific assay composed primarily of a primer pair encompassing a region in the context of the CpG site of interest, containing at least one recognition site for methylation-sensitive restriction enzymes. The second pair of primers targets a DNA region away from any CpG island and whose amplimer does not include recognition sites for methylation-sensitive restriction enzymes (used as restriction enzyme-resistant control amplimer). The third pair of primers targets a CpG site overlapping a methylation-sensitive restriction-enzyme recognition site located in the *ESCO2* core promoter CpG island in chromosome 8, and which is known to be consistently unmethylated in multiple human tissues (used as restriction enzyme-susceptible control amplimer) (Alves da Silva et al., 2016). The forward primers were labeled with a fluorochrome for visualization, and the amplimers were analyzed and quantified comparatively by capillary electrophoresis in an automated laser fluorescent ABI PRISM 310 Genetic Analyzer (Thermo Fisher Scientific, Waltham, MA, USA). The electropherograms were produced with the GeneScan® Analysis and Genotyper® software version 3.7 packages and GeneMapper® ID version 3.2 (Applied Biosystems ® from Thermo Fisher Scientific, Waltham, MA, USA). The level of methylation at the individual CpG sites was estimated as the normalized ratio of restriction enzyme-resistant amplimer after digestion as compared with the undigested product and corrected by the proportion of resistant positive control amplimer, using the formula previously reported (Alves da Silva et al., 2016; de Sa Machado Araujo et al., 2018). Target amplimers ranged 150 bp to 440 bp in length Control amplimers (resistant to enzymatic digestion) were ten bp larger than the target amplimers (Supplemental Table S3).

### Characterization of Alleles

To the amplification products were added formamide (Hi-Di Formamide, Applied Biosystems) and the GeneScan 500 molecular weight standard labeled with the fluorochrome LIZ ™ (orange fluorescence) (Applied Biosystems). The amplification products were separated by high resolution ((1 bp) capillary electrophoresis in the POP4 polymer using the ABI Prism ™ 310 Genetic Analyzer platform. The electrophoretic profiles were analyzed using the GeneMapper ID 3.2 programs (Applied Biosystems).

### Computational Cross-Referencing with 5meCpG Levels in Public Methylomes

We validated the levels of methylation at each specific CpG site in the target immune regulatory/response genes observed in healthy individuals by computationally cross-referencing with data values from public genome-wide methylomes (bisulfite sequencing - BS-Seq - experiments). We extracted data from the integrative analysis of 111 reference epigenomes (Roadmap Epigenomics et al., 2015), available from the University of California Santa Cruz (UCSC) Genome Browser (Kent et al., 2002; Raney et al., 2014). We selected BS-Seq experiments performed in DNA from whole-blood (newborn and 100-year old donors), and the following blood-derived enriched cell types: B-cell, T-cell, NK-cells, neutrophils, and macrophages. We also included the methylomes from liver, lung, spleen, and thymus. Details of the methylome studies are provided in Supplemental Table S4 Dataset A. The methylation values were extracted using UCSC Genome Browser Data Integrator web tool (Hinrichs et al., 2016) and the 5meCpG site values were plotted using the *ggplot2* graph data analysis (Wickham, 2016) in R-codes (R Core Team, 2018).

### Cross-Referencing with (Epi)genotype-Phenotype Associations

We lookup online public database about GWAS studies for evidence on the SNP and CpG sites interrogated to identify possible phenotypes and disease-associated loci or expression quantitative trait loci (eQTLs) and to assign chromatin states to the lead variants and CpG sites. The strategy was as described previously for the differentially methylated regions associated with genomic imprinting (Machado et al., 2014). Briefly, the SNP rsID# or the physical coordinates were confronted for evidence in the Phenotype-Genotype Integrator - PheGen*I* (Ramos et al., 2014), e-GRASP (Karim et al., 2016), PhenoScanner (Staley et al., 2016), and HaploReg v4.1 (Ward and Kellis, 2016) web database tools. We restricted the search to *p*-and r2 cut-off values ≥ 5 × 10-8 and 0.8, respectively, to warrant evidence with genome-wide scores.

### Statistical Analysis

The methylation levels at the individual CpG sites were evaluated for normal distribution using the Shapiro-Wilk test of normality with R-code script (R Core Team, 2018). When the pair of groups presented methylation levels at the same individual CpG sites with a normal distribution, the groups were compared using the Welch’s *t*-test (unequal variances *t*-test). Otherwise, the groups were analyzed using the Wilcoxon Rank Sum Test to calculate *p*-values and 95% confidence intervals (95%CI) of the difference between the different groups. The range of 95%CI was graphically displayed as a forest plot using *ggplot2* graph data analysis.

## Results

To select the target genes, we consider the immune regulatory mechanism of production of anti-FVIII inhibitor antibodies. The exact mechanism of inhibitor production has not yet been clarified, but it is proposed to initiate when antigen-presenting cells (APC) endocytose the exogenous FVIII and degrade it into peptides. In turn, these peptides bind to molecules of the major histocompatibility complex (HLA) class II, and presented to CD4 + T-cells in the secondary lymphoid organs (Figure 1) and can receive CD28 stimulatory signals, or CTLA4 inhibitory signals. Additionally, small fragments of the endogenous FVIII can be produced by the patient and presented via HLA class I inducing activation and clonal expansion of CD8 + T-cells. In this form, CD4 + T lymphocyte clones induce B lymphocytes to produce autoantibodies through the interaction of CD40 receptors with CD40L. The induction of lymphocytes is dependent on cytokines and other molecules such as IL1B, IL2, IL6 and CST7. In humans, both Th_1_ and Th_2_ responses induce the synthesis of anti-FVIII antibodies in hemophiliac patients. However, there is a predominance of Th_2_ response producing IgG4 antibodies (Reding et al., 2002; Chaves et al., 2010b). The polarized response may be directed primarily by cytokines such as IL4, IFN-γ, and TNF-α. The increased immune response, whether inflammatory or anti-inflammatory, may induce IL10 regulatory cytokine production.

**FIGURE 1.**
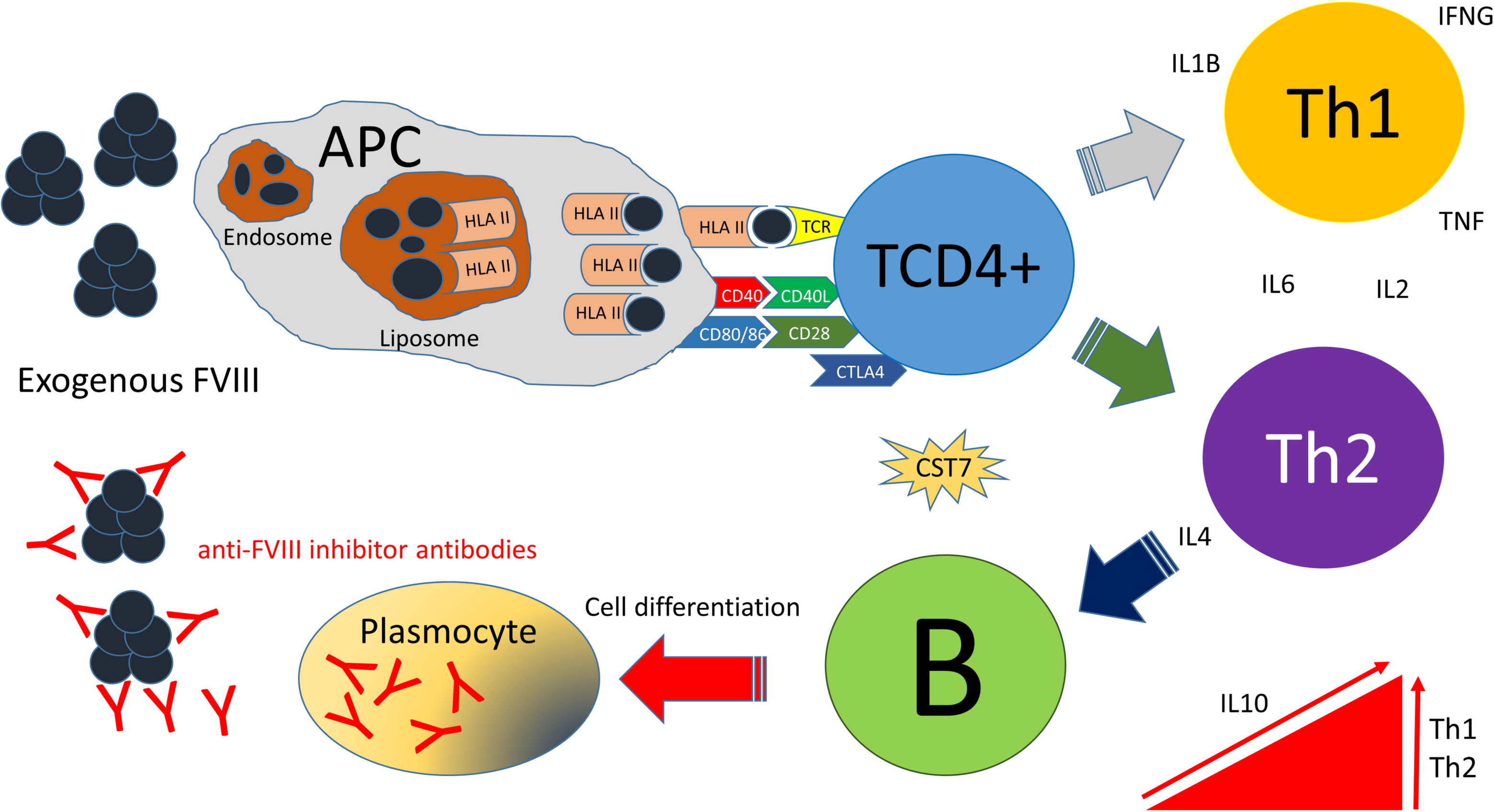
Cartoon representation of the proposed immune regulatory mechanisms linked to the development of anti-FVIII inhibitor antibodies in hemophiliac patients undergoing FVIII replacement therapy. Details are given in the main text.

Thus, our criteria to include an immune regulatory/ response gene included: (*i*) evidence of regulation of the promoter region by DNA methylation (Table 1); (*ii*) occurrence of allele variants reported in association with the risk of inhibitor development (*IL1B, IL6, IL10, TNF* and *CTLA*) (Astermark et al., 2006a; Astermark et al., 2006b; Astermark et al., 2007; Lozier et al., 2011; Repesse et al., 2013); (*iii*) genes with differential expression in patients presenting with inhibitors (*CST7*) (Santos, 2010; Hwang et al., 2012); (*iv*) cytokine coding genes whose protein dosage presents alteration in individuals with inhibitors (*IFNG* and *IL4*) (Oliveira et al., 2013); (*v*) genes with modulator function in the immune response mediated by antibodies or innate immunity (*IL2, CD28*). A summary of research articles about methylation in the target genes is presented in Table 2.

**Table 1.** Genes selected from the literature due to their association with the development of anti-FVIII inhibitor antibodies and the regulation of promoter regions by DNA methylation.

| Gene | Association with inhibitor development | Promoter regulation by CpG methylation |
| --- | --- | --- |
| *CTLA4* | (Astermark et al., 2007) | (Cribbs et al., 2014) |
| *IFNG* | (Hu et al., 2007) | (White et al., 2002; Kwon et al., 2008) |
| *IL10* | (Astermark et al., 2006b) | (Szalmas et al., 2008) |
| *IL1B* | (Lozier et al., 2011) | (Hashimoto et al., 2009) |
| *IL2* | (Lozier et al., 2011) | (Murayama et al., 2006) |
| *IL4* | (Hu et al., 2007) | (Kwon et al., 2008; Mi and Zeng, 2008) |
| *TNF* | (Astermark et al., 2006a) | (Campion et al., 2009; Gowers et al., 2011; Hermsdorff et al., 2013) |
| *IL8* | (Hwang et al., 2012) | (Dimberg et al., 2011) |
| *IL6* | (de Alencar et al., 2015) | (Nile et al., 2008) |

**Table 2.** Available published data on DNA methylation in the target genes.

| Reference | Gene | Function | Methylation |  | Cell line/source | Findings |
| --- | --- | --- | --- | --- | --- | --- |
|  |  |  | Position | Status |  |  |
| (Cribbs et al., 2014) | *CTLA4* | Signaling | Promoter | Hyper | Treg | Treg promoter methylation in patients with rheumatoid arthritis |
| (White et al., 2002) | *IFNG* | Inflammation | Promoter | Hyper | Umbilical cord CD4+ T-cells | Promoter hypermethylation in umbilical cord CD4 + T lymphocytes compared to adult cells, which is associated with lower *IFNG* expression in newborns compared with adult CD4 + T-cells |
| (Szalmas et al., 2008) | *IL10* | Antibody production | Promoter | Hyper | Cervical cancer cells | Hypermethylated promoter region in cervical cancer cell lines |
| (Hashimoto et al., 2009) | *IL1B* | Inflammation | Promoter | Hypo | Human chondrocyte | Increased gene expression with the administration of AZA and IL1B. There is a reduction in promoter methylation |
| (Murayama et al., 2006) | *IL2* | Cell proliferation | Promoter | Hypo | CD4 T-cells | Loss of methylation in a single CpG site increases expression |
| (Kwon et al., 2008) | *IL4* | Inflammation | Promoter | Hypo | CD4 T-cells | After antigenic stimulation with *Dermatophagoides*, methylation decreases in CD4 + T-cells in patients with bronchial asthma. The concentration of IL4 showed a correlation with the level of methylation |
| (Nile et al., 2008) | *IL6* | Cell proliferation | Promoter | Hypo | PBMC | Some hypomethylated CpG sites in patients with rheumatoid arthritis. Methylation correlates with expression |
| (Campion et al., 2009) | *TNF* | Inflammation | Promoter | Hypo | PBMC | Obese men with weight loss on a calorie restriction diet have loss of methylation at the *TNF* promoter |

We determined the methylation status of thirteen individual CpG sites in ten target genes (Supplemental Table S3) using restriction enzyme methylation-sensitive PCR triplex assays. For each CpG site, a specific MSRE-PCR assay was developed. We required at least one methylation-sensitive restriction enzyme recognition site overlapping the selected CpG site within the specific amplimer. Figure 2 illustrates the genomic context of the three CpG sites (CpG-1, CpG-2, CpG-3) profiled in the *IL4* gene. The genomic context of the individual CpG sites for the other genes is illustrated in Supplemental Figures S1-S9.

**FIGURE 2.**
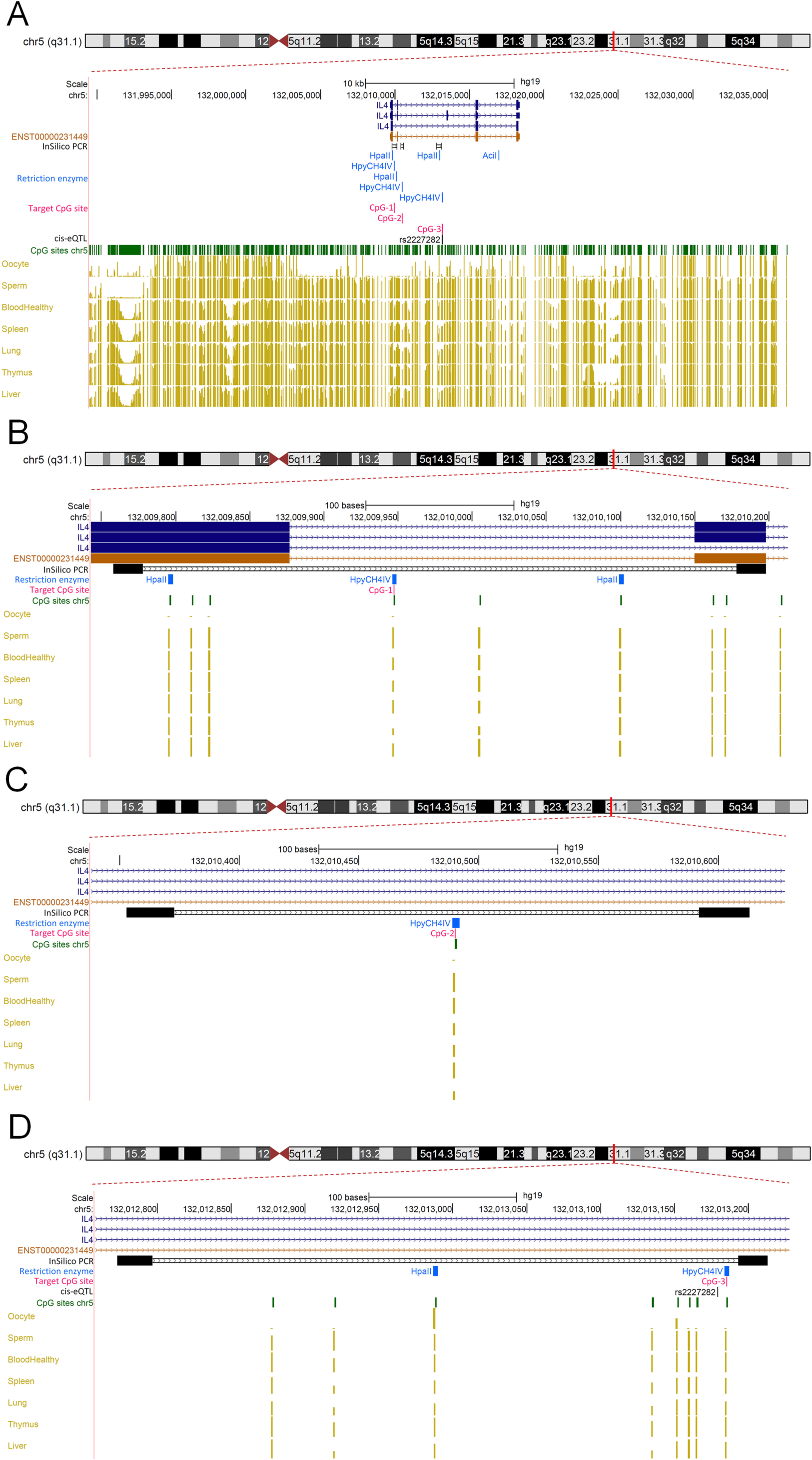
The *IL4* locus and the associated CpG sites interrogated. (**A**) Screenshot across a 40-kb long-span view (hg19; chr5:131994661-132034660) centered at the *IL4* gene locus. The annotated features are (from top to bottom): chromosome 5 ideogram; physical positions and the exon-intron organization of the reference and alternative *IL4* transcript isoforms (dark blue color), which are transcribed in the forward direction from the plus DNA strand; the *IL4* APPRIS principal isoform ENST00000231449 (brown color) (Rodriguez et al., 2013); the relative positions of the three in silico generated amplimers (black color) encompassing the CpG sites individually interrogated in the present study (pink color); the relative position of the *cis*-eQTL SNP rs2227282 (black color); the relative positions of the restriction enzyme sites susceptible to DNA methylation (light blue color ticks); the overall CpG sites across the region (light green ticks); the methylation status at the CpG sites (golden ticks) across a 40-kb long-span view reported in public methylomes from gametes (oocyte and sperm) and somatic tissues (blood, spleen, lung, thymus and liver). The methylation levels are represented on a scale from 0 to 1 (hypomethylated to hypermethylated). Note the overall hypomethylation statuses of the CpGs spanning the *IL4* gene in DNA from oocytes in contrast to the hypermethylated statuses in sperm and somatic tissues. Therefore, in the *IL4* gene region, there is asymmetrical methylation in gametes (hypomethylation in oocytes and hypermethylation in spermatozoa). Screenshot prepared using public and custom hubs available from the UCSC Genome Browser (Kent et al., 2002; Raney et al., 2014). (**B**) Zoom-in screenshot across a 460-bp long-span view (hg19; chr5:132009759-132010198), encompassing the 440bp long amplimer that contains the *IL4* CpG-1 site interrogated in the study. The annotated features are as in panel (**A**). (**C**) Zoom-in screenshot across a 300-bp long-span view (hg19; chr5:132010354-132010613), encompassing the 260bp long amplimer that contains the *IL4* CpG-2 site interrogated in the study. The annotated features are as in panel (**A**). (**D**) Zoom-in screenshot across a 460-bp long-span view (hg19; chr5:132012774-132013213), encompassing the 440bp long amplimer that contains the *IL4* CpG-3 site interrogated in the study. The annotated features are as in panel (**A**).

Initially, we interrogated all the thirteen specific CpG sites in a subset of samples (up to ten cases and ten controls) and ten non-hemophilia A healthy subjects. At the particular CpG sites, we consistently observed one of three possible methylation statuses: hypermethylated (>99% restriction enzyme-resistant amplimers), hypomethylated (<1% resistant) and intermediate methylated (close to 50% resistant). We observed a hypermethylated status at the individual CpG sites at *CD28, IFNG, CTLA4* (CpG-1), *CTLA4* (CpG-2), *IL4* (CpG-2), and *IL4* (CpG-3). The selected CpG sites at the *IL2, IL1B* and *TNF* genes were hypomethylated. The selected CpG sites at the *IL4* (CpG-1), *IL10, CST7*, and *IL6* genes exhibited an intermediate methylation status. Representative examples of the MSRE-PCR assays are in Supplemental Figure S10. The distribution of methylation levels was similar among the three groups (Figure S11). In the discovery set limited to ten individuals per group, no significant statistical difference was found for any of the targets between the three groups of subjects (p>0.05). Based on the observed profiles and the extent of the interindividual variation in methylation levels, we narrowed down the analysis to five specific CpG sites to profile the full set of twenty patients with inhibitors, twenty patients without inhibitors and twenty non-hemophilia A subjects. The three targets in the *IL4* gene were selected, one in *IL6* and one in *CST7*. The *IL4* and *IL6* CpG site were reappraised based on the role of interleukin B lymphocytes proliferation and the CpG site at the *CST7* because of evidence of *CST7* altered expression in hemophilia A patients with inhibitors (Santos, 2010). Although the distribution of methylation levels was similar among the three groups (Figure 3), we observed a discrete, yet statistically significant decrease in the methylation levels at the *IL4* (CpG-3) between the affected patients presenting with anti-FVIII inhibitors as compared with those without (95% CI [-1.009e-01-5.149e-5]), Wilcoxon Rank Sum Test with continuity correction *p* = 0.04; Figure 4 and Supplemental Table S5). We also noted that all the evaluated CpG sites at the target genes showed concordant levels of methylation reported in several methylome studies from diverse biological samples withdrawn from healthy subjects (Supplemental Table S4 Dataset B).

**FIGURE 3.**
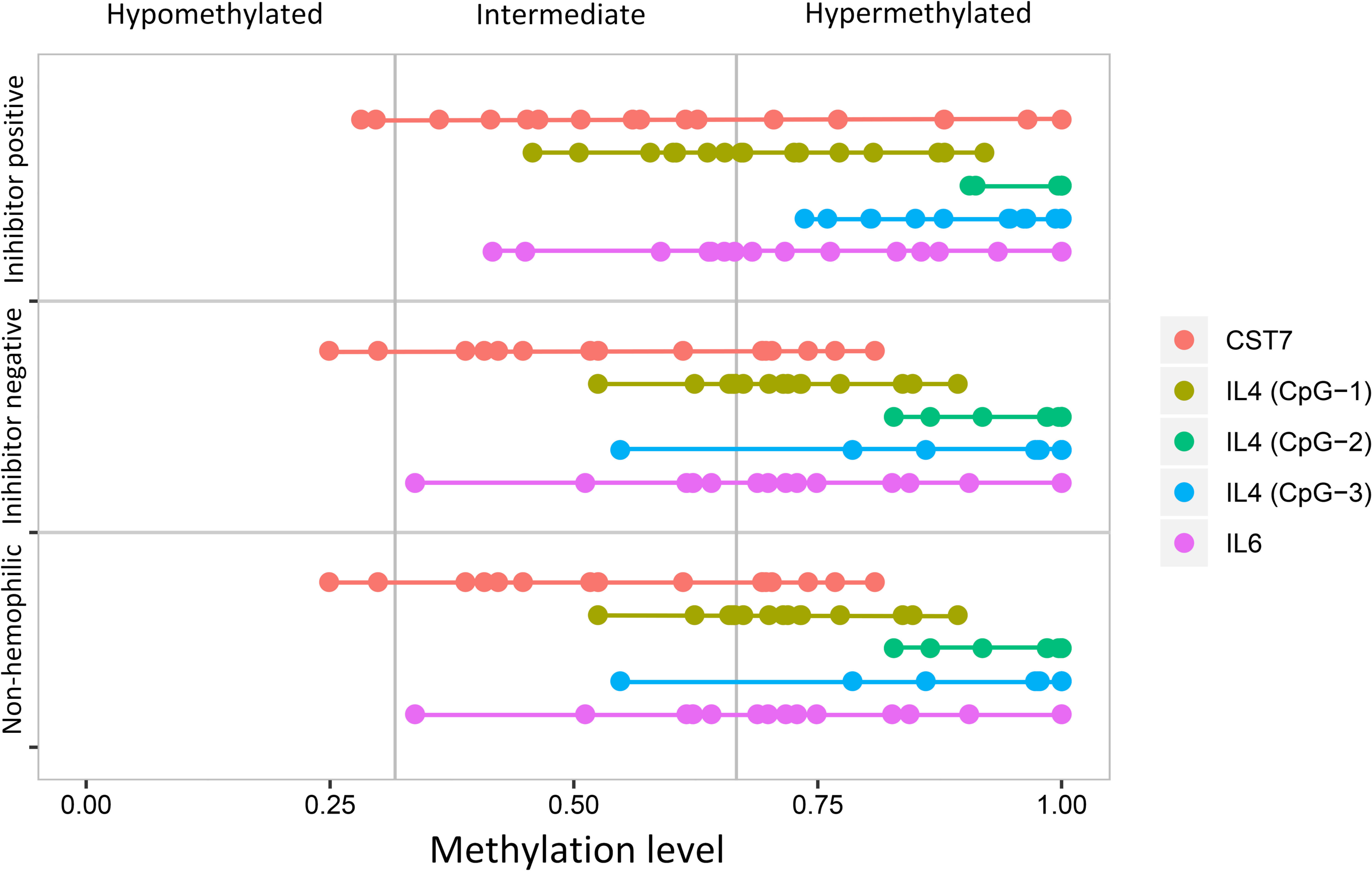
The range of the methylation levels at the individual CpG sites in the *IL4, Il6*, and *CST7* genes. The methylation statuses are classified in three possible patterns: hypermethylated, intermediate methylated and hypomethylated. The three different groups of subjects (n = 20 per group) are shown on the left. Names of the individual CpG sites are displayed on the right, and each site is depicted in a different color.

**FIGURE 4.**
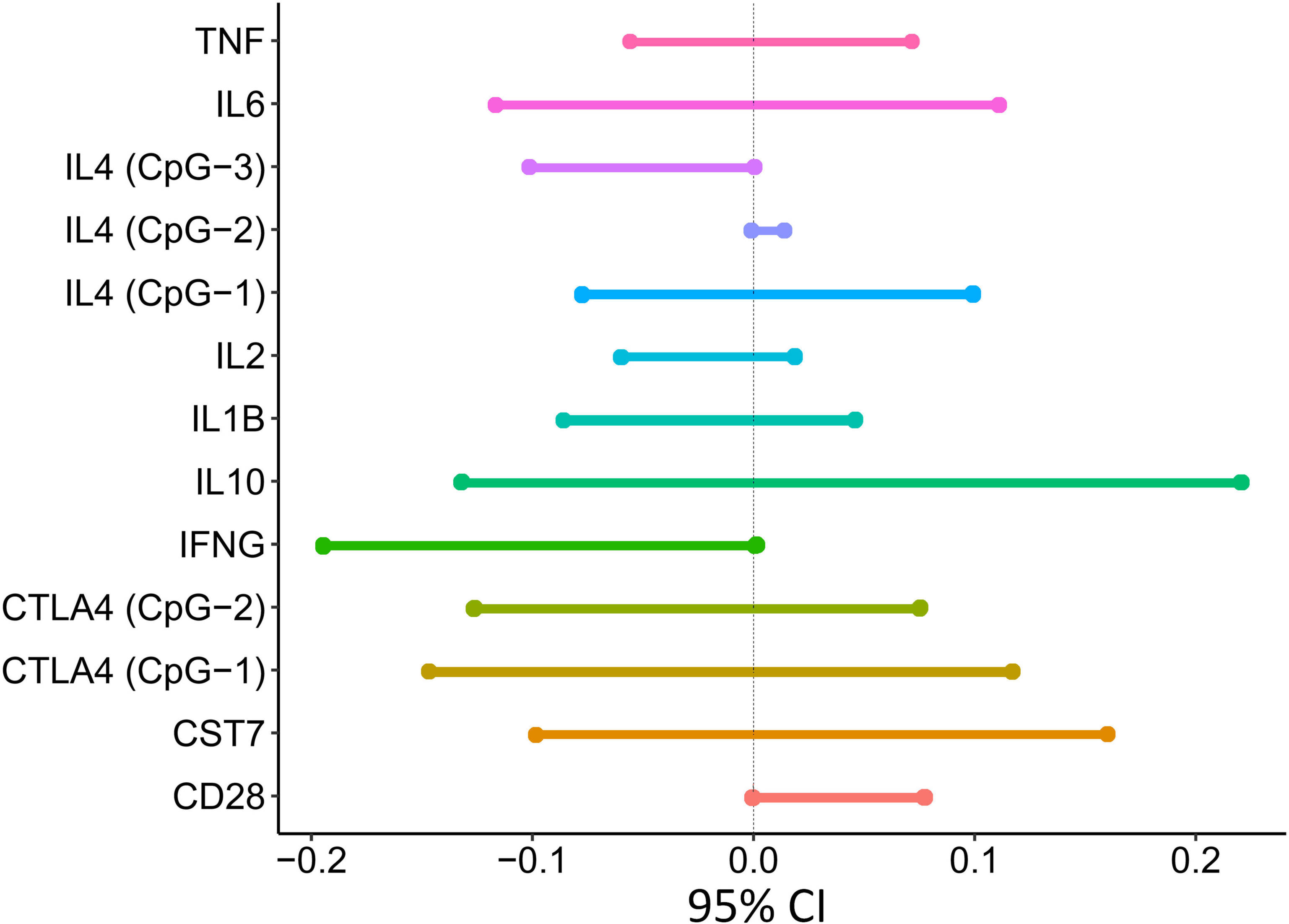
Forest plot showing the association between inhibitor development and levels of methylation at five individual CpG sites. Graphic display of the associated confidence intervals from the comparison of methylation states at five individual CpG sites between the groups of hemophilia A patients who developed anti-FVIII inhibitor antibodies and those who did not produce them. The x-axis forms the effect size scale (95% CI), which represents the estimate of an interval (horizontal lines) in which the “true” effect (in the population) will most probably lie. The vertical dashed line represents the meta-analysis summary measure. Names of the individual CpG sites are shown on the left.

Within the *IL6* amplimer, there is a two-base pair deletion, referring to the SNP rs35081782 (NC_000007.13:g.22765338_22765339CT[3]). Although the deletion had a lower frequency in the group of hemophilia A with inhibitors (15%), no significant difference was found in the distribution of the variant between the three groups here evaluated (Table S6). Cross-referencing with the PhenoScanner GWAS public database, we found that in one study (Astle et al., 2016) the rs35081782 indel variant has been associated at genome-wide scores with white blood cell count traits linked to common complex diseases (Table S7 Dataset A).

Cross-referencing with 5meCpG values from seven public data BS-Seq methylomes (Supplemental Table S4 Dataset A) corroborated the overall methylation profiles generated by MSRE-PCR. Hypermethylated, hypomethylated and intermediately methylated patterns were consistent with those observed in blood-derived cell types and other tissues (Figures 5 and 6 and Supplemental Table S4 Dataset B).

**FIGURE 5.**
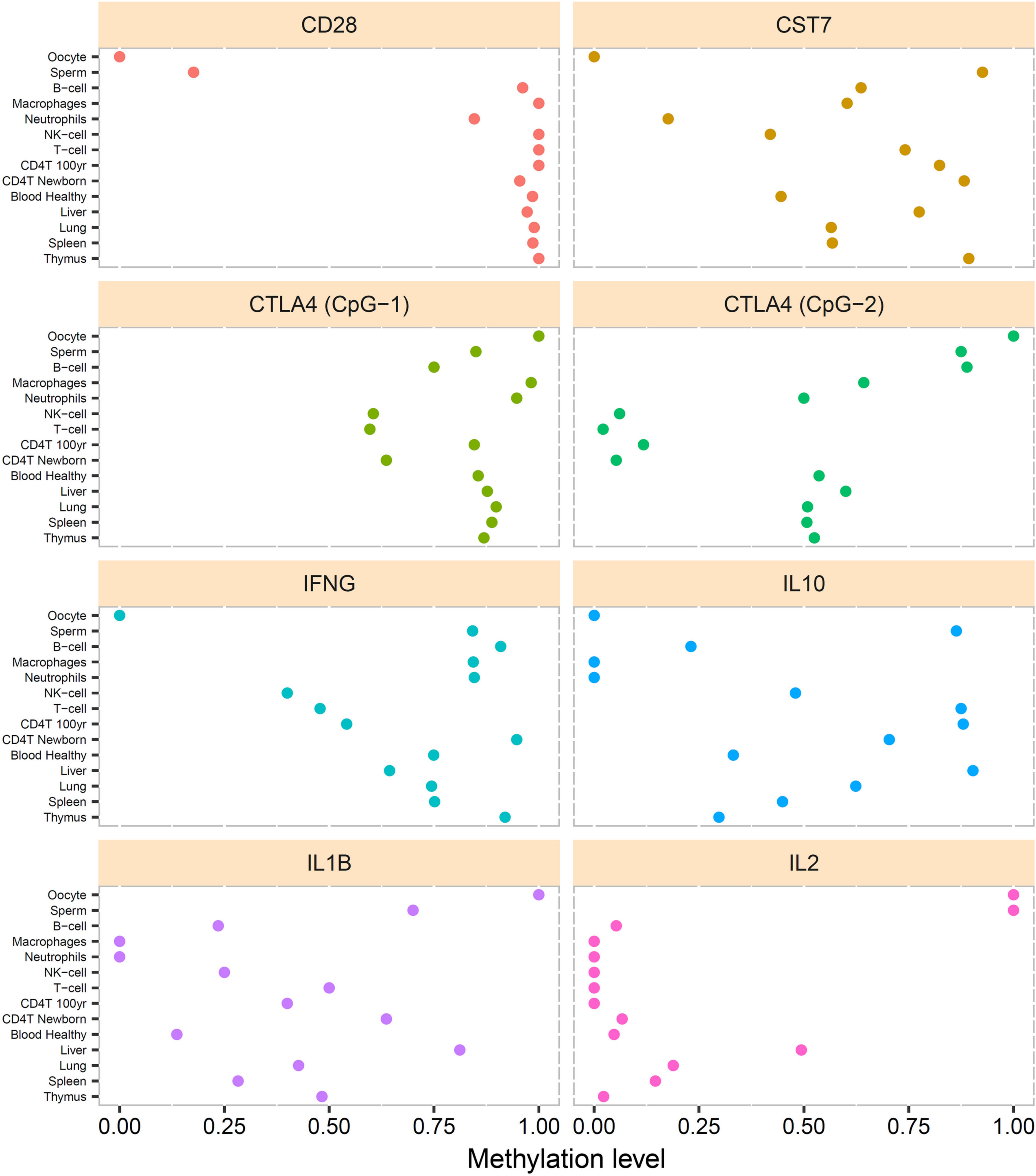
Distribution of the methylation statuses at eight individual CpG sites in public methylomes. The x-axis represents the extent of methylation at the individual CpG sites in the *CD28, CST7, CTLA4, IFNG, IL10, IL1B*, and *IL2* genes (shown on the top of each panel). The methylome studies (see Table S4 Dataset (**B**) for details) are named on the left according to the blood-derived cell type, whole blood or gametes.

**FIGURE 6.**
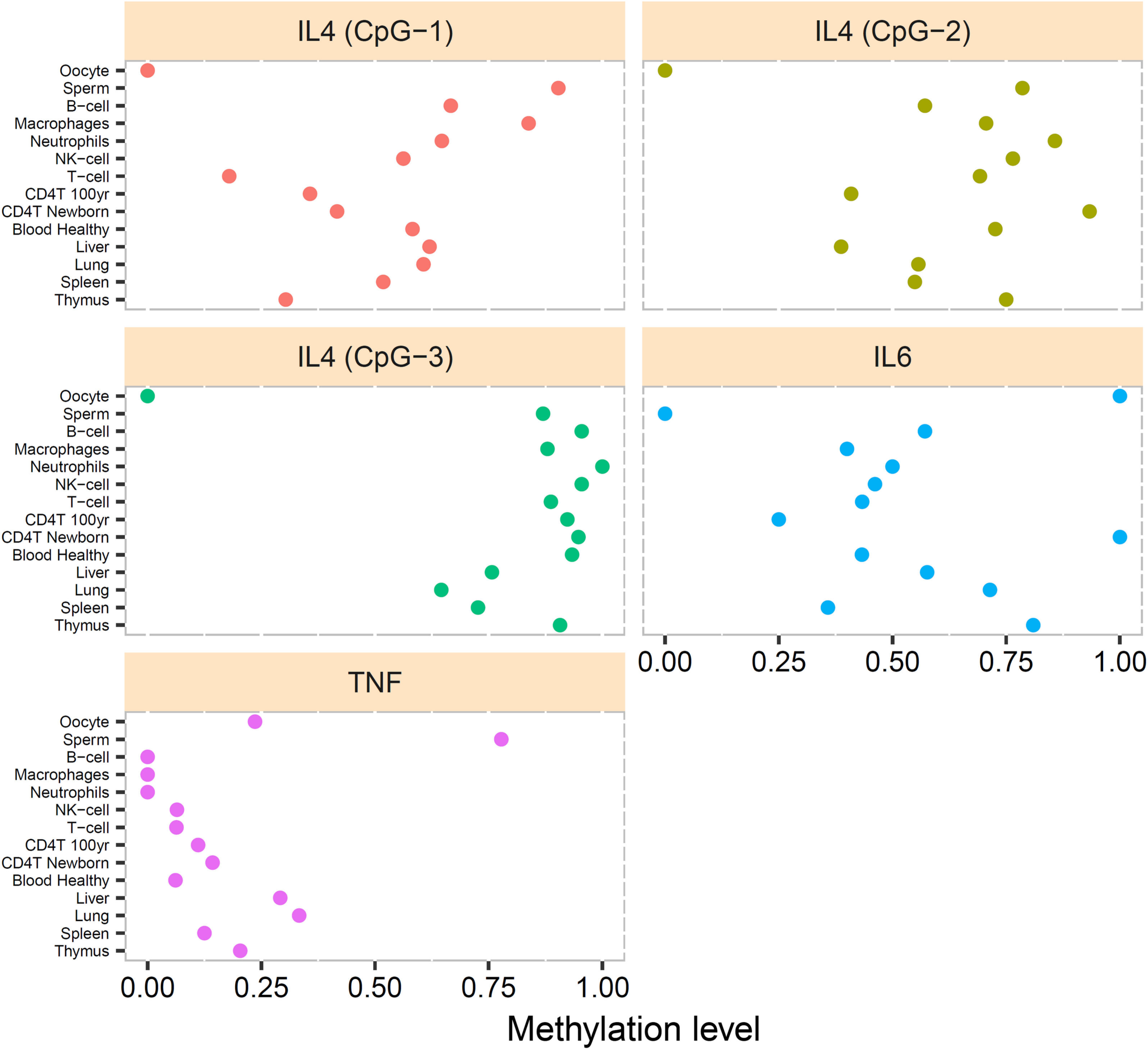
Distribution of the methylation statuses at five individual CpG sites in public methylomes. The x-axis represents the extent of methylation at the individual CpG sites in the *IL4, IL6* and *TNF* genes (shown on the top of each panel). The methylome studies (see Table S4 Dataset (**B**) for details) are named on the left according to the blood-derived cell type, whole blood or gametes.

We also noted that most of the 5meCpG sites (*IL10, IFNG, CST7, IL4*, and *TNF*) exhibited gametic asymmetry in that the oocyte’s DNA was hypomethylated and the sperm’s DNA was hypermethylated (Table S4 Dataset B). The gametic-of-origin asymmetry was not maintained in adult tissues since the level of methylation shifted to a hypermethylated or a hypomethylated state.

By cross-referencing the physical coordinates of the region containing the *IL4* CpG-3 site (chr5:132013135-132013234) with data from the integrative analysis of 111 reference epigenomes (Roadmap Epigenomics et al., 2015), we noted that the differentially methylated CpG-3 maps within an enhancer domain, which overlaps a DNase susceptibility region in intron 1 of the *IL4* gene. The enhancer domain is enriched with the H3K27ac and H3K4me1 histone modification active marks in chromatin from several tissues but to a much lesser extent in blood or blood-derived cells types (Supplemental Table S7 Dataset B). This chromatin configuration is, therefore, permissive for gene expression. Within the region encompassing the lead *IL4* 5meCpG site multiple 5′-cap active transcriptional start sites (CAGE-TSS) have been detected (Severin et al., 2014; Lizio et al., 2017).

We also noted the occurrence of the SNP rs2227282 (NC_000005.9:g.132013179C>G) 6-bp upstream of the *IL4* (CpG-3) site, and which acts as an expression quantitative trait locus (eQTL) affecting the transcript levels of the *SLC22A4* and *SLC22A5* genes in peripheral blood monocytes (Zeller et al., 2010). rs2227282 also influences the RNA expression levels of the *IL4* (Westra et al., 2013), *KIF3A, SEPT8*, and *PDLIM4* (Lappalainen et al., 2013; The GTEx Project, 2015), and *RAD50* (Westra et al., 2013) genes (Table S7 Dataset C). In GWAS, rs2227282 has been associated with high confidence scores to allergic diseases (Table S7 Dataset A).

Using the STRING v11 database of protein-protein association networks (Szklarczyk et al., 2018), we found no high confidence network of interactions between the protein products of the genes whose transcript expression levels are affected by the *IL4* intronic rs2227282 *cis*-eQTL (Table S7 Dataset D).

## Discussion

Although associations between genetic factors and the production of anti-FVIII inhibitor antibodies in hemophilia A have been previously investigated, the investigation about the contribution of epigenetic marks is lacking. Our study is the first to report a differentially methylated CpG epigenetic mark between people who develop and do not develop inhibitors upon replacement therapy, characterized by a statistically significant decreased methylation level in the group of patients presenting with inhibitors. The differentially methylated CpG site is located within an enhancer domain which overlaps a DNase hypersensitive region in the intron 1 of the *IL4* gene. Six bp upstream of the *IL4* 5meCpG-3 site, there is the rs2227282 SNP that acts as *cis*-eQTL by influencing the transcript levels of the *PDLIM4, SLC22A4, SLC22A5, RAD50, IL4, KIF3A, SEPT8*. These six genes are physically linked (ordered as shown above) within a 520 kb domain in chromosome 5 that encompasses the Th_2_ *IL5, IL13*, and *IL4* cytokine gene cluster locus. The protein products encoded by the seven genes influenced by the rs2227282 *cis*-eQTL are not predicted to directly interact in a functional network with high confidence of enrichment in the following categories: Cellular Component (Gene Ontology), KEGG Pathways, PFAM Protein domains, INTERPRO protein domains and features (Table S7 Dataset D).

Although two previous studies investigated epigenetic 5meCpG changes in the *F8* gene, they were not in the context of inhibitor’s development. The first study (El-Maarri et al., 1998) explored the fact that transitional mutations at CpG dinucleotides account for approximately a third of all germline point mutations, which arise through spontaneous deamination of 5meCpG as a sort of causative point mutation in index patients with unknown mutations. Using *F8* gene-based BS-Seq, the authors screened for such mutations occurring at the *F8* gene in mature male and female germ cells. The *F8* gene was found to be equally and profoundly methylated in oocytes and spermatocytes. In the second study in eighty hemophilia A male index patients with unknown mutation status and no information about anti-FVIII inhibitor antibodies (Zimmermann et al., 2013), blood DNA samples were screened for epigenetic changes by bisulfite pyrosequencing assays in the 5’UTR and intron 1 regions of the *F8* gene. The methylation patterns observed in the hemophilia A patients did not differ from those in the controls, with the 5’UTR CpG islands being consistently hypomethylated whereas CpG islands at the intron 1 being systematically hypermethylated.

Our analysis indicates that only the *IL4* (CpG-3) site exhibited statistical differences at the methylation statuses between the group of hemophilia A patients who are positive for inhibitors and the group of patients who are negative for the anti-FVIII antibodies. We envision the *IL4* (CpG-3) site as a promising lead epigenetic mark, whose potential value must be appraised in a more significant number of patients. IL-4 has immunomodulatory activity, being one of the main effectors in the polarization for a Th_2_ response. As pointed earlier, the *IL4* gene is located within the Th_2_ immune response regulatory *IL5, IL13*, and *IL4* cytokine locus. The Th_2_ region has several sites for binding of the chromatin remodeling transcription factor GATA3, which stimulates the Th_2_ locus to express several genes within the Th_2_ locus (Wilson et al., 2009; Adkins and Yoshimoto, 2014). It also has several sites for T-BET binding, a transcription factor that represses expression (Kanhere et al., 2012). Since the anti-FVIII inhibitor antibodies belong mainly to the IgG4 subclass (Montalvao et al., 2015), a Th_2_ response may be more pronounced in individuals who develop inhibitors (Reding et al., 2002; Chaves et al., 2010a; Chaves et al., 2010b; Oliveira et al., 2013). Future assessment of the correlation between the *IL4* differentially methylated (CpG-3) site and IL-4 production must consider all aspects mentioned above.

Some studies have found that both Th_1_ and Th_2_ responses are pronounced in patients who are positive for inhibitor antibodies (Hu et al., 2007). Th_1_ cells produce the inflammatory cytokine IFN-γ. It is known that methylation of the *IFNG* gene promoter regulates its expression (White et al., 2002). Therefore, we also expected differences in the methylation level at the CpG site of the *IFNG* gene. Our data showed a lower mean level of methylation in the group presenting with inhibitors, but the differences between the methylation statuses at the individual CpG site interrogated in the *IFNG* gene had a *p*-value of 0.061 (Table S5). Whether the observed differences result in measurable increments in the production of IFN-γ is unclear.

We note two potential caveats against the results of our study. Firstly, the limited number of twenty subjects per group may have impaired detecting more subtle differences among groups. Secondly, the testing of whole blood DNA may mask discrete differences in methylation levels specific to some cell populations. Regarding the sample size, we included the maximum number of subjects available with the same class of causative mutation (Inv22). Moreover, all the hemophilia A patients were from the same hemophilia center in Argentina to minimize possible confounding effects that are secondary to different ethnicities or admixed population, as the Brazilian population. Moreover, most of the analyzed samples came from patients bearing the Inv22-1 mutation (n = 34; where n = 17 with inhibitors and n = 17 without). The other six samples came from patients who had the Inv22-2 mutation (n=3 with and n=3 without inhibitors). Perhaps through an international consortium, a much broader set of samples can be included to reappraise statistical significance and to enable the identification of other lead CpG sites, ideally, using a genome-wide methylome (BS-Seq) strategy. Unfortunately, no such methylome studies have been carried out in hemophilia A patients presenting with inhibitors. Our results also point to extensive interindividual variation in methylation levels at the selected CpG sites, indicating that the statuses of the selected sites are affected by, as yet, unknown determinants.

Regarding the second caveat, several genome-wide methylome studies in blood samples from healthy individuals have reported that the methylation levels at individual CpG sites can reflect the apparent differences in the distribution of cell-type populations in whole blood (Houseman et al., 2012; Jaffe and Irizarry, 2014). There are CpG sites that exhibit cell type-dependent methylation, being either hypermethylated or hypomethylated in different blood-derived cell types (i.e., B-cells, NK cells) (Figure S12, and Supplemental Table S4 Dataset B). Regarding the possible confounders due to varying densities of cell types in whole blood, the argument would gain in strength if significant differences had been found for every sample among the groups here tested. Nevertheless, through computational cross-referencing with methylome studies in whole blood and blood-derived cell fractions from healthy individuals, we estimated that the methylation statuses of the CpG sites here interrogated were concordant with the profiles obtained by the indirect MSRE PCR assays. Thus, the CpG sites examined might not be subject to the cell density caveat mentioned above.

The methylation statuses at CpG sites can also be affected by gender (McCarthy et al., 2014; Lee et al., 2018) and age (Lin et al., 2016; Zhang et al., 2017). A gender-related effect is ruled out in our experimental setting because the patients were males. However, it is possible that part of the variation of the level of methylation observed in the individual sites here interrogated is related to differences in the age of the subjects at the collection time. In the subset of patients with inhibitors, the mean age was 12.4 years whereas in the group without inhibitors was 13.85 years. Noteworthy, in the group of healthy individuals the mean age was 52.3 years. By cross-referencing with methylation levels for public methylomes from healthy donors, we noted that most CpG sites tested were equally methylated at extreme ages, including 100-year old individuals, and whether being hypomethylated, intermediate methylated or hypermethylated, with the exception of the CpG sites at the *IL4* (CpG-2) and *IL6* genes, that tend to be hypermethylated in newborn but intermediately methylated at very advanced age. Since no significant difference was observed in the methylation levels after age stratification, we concluded that the differences in methylation levels at most CpG sites interrogated in the present study are not age-related.

## Web Resources

The URLs for data presented herein are as follows:

UCSC Genome Browser, https://genome.ucsc.edu/

Roadmap Epigenomics Browser, http://www.roadmapepigenomics.org/

WashU Epigenome Browser, http://epigenomegateway.wustl.edu/

FANTOM ZENBU Browser, http://fantom.gsc.riken.jp/zenbu/

PhenoScanner. http://www.phenoscanner.medschl.cam.ac.uk/phenoscanner

e-GRASP, http://www.mypeg.info/egrasp

HaploReg, http://compbio.mit.edu/HaploReg

PheGen*I*; https://www.ncbi.nlm.nih.gov/gap/phegeni

R software package, http://www.R-project.org

## Author Contributions

TBS, EM-A: designed the study, analyzed data, wrote the manuscript. TBS, TLS: performed MSRE-PCR typing and performed statistical analysis. LCR, VDM, CDDB: set up the cohort of hemophilia A patients, performed inverse sighted-PCR and created the clinical databank. CSF, CFOS, EM-A: carried out cross-referencing with public epigenomic studies and prepared figures. All the authors gave final approval.

## Acknowledgments

We thank all participants and their guardians for the superb willingness to contribute to this study.

## Supplemental Materials

**FIGURE S1. The *CD28* locus and the associated CpG sites interrogated**. (**A**) Screenshot across a 40-kb long-span view (hg19; chr2:204591638-204592077) centered at the *CD28* gene locus. The annotated features are (from top to bottom): chromosome 2 ideogram; physical positions and the exon-intron organization of the reference and alternative *CD28* transcript isoforms (dark blue color), which are transcribed in the forward direction from the plus DNA strand; the *CD28* APPRIS principal isoform ENST00000324106 (brown color) (Rodriguez et al., 2013); the relative positions of the three in silico generated amplimers (black color) encompassing the CpG sites individually interrogated in the present study (pink color); the relative positions of the restriction enzyme sites susceptible to DNA methylation (light blue color ticks); the overall CpG sites across the region (light green ticks); the methylation status at the CpG sites (golden ticks) across a 40-kb long-span view reported in public methylomes from gametes (oocyte and sperm) and somatic tissues (blood, spleen, lung, thymus and liver). The methylation levels are represented on a scale from 0 to 1 (hypomethylated to hypermethylated). Note the overall hypomethylation statuses of the CpGs spanning the *CD28* gene in DNA from oocytes in contrast to the hypermethylated statuses in sperm and somatic tissues. Therefore, in the *CD28* gene region, there is asymmetrical methylation in gametes (hypomethylation in oocytes and hypermethylation in spermatozoa). Screenshot prepared using public and custom hubs available from the UCSC Genome Browser (Kent et al., 2002; Raney et al., 2014). (**B**) Zoom-in screenshot across a 460-bp long-span view (hg19; chr2:204591638-204592077), encompassing the 440bp long amplimer that contains the *CD28* CpG-1 site interrogated in the study. The annotated features are as in panel (**A**)

**FIGURE S2. The *CTLA4* locus and the associated CpG sites interrogated**. (**A**) Screenshot across a 40-kb long-span view (hg19; chr2:204714572-204754650) centered at the *CTLA4* gene locus. The annotated features are (from top to bottom): chromosome 2 ideogram; physical positions and the exon-intron organization of the reference and alternative *CTLA4* transcript isoforms (dark blue color), which are transcribed in the forward direction from the plus DNA strand; the *CTLA4* APPRIS principal isoform ENST00000302823 (brown color) (Rodriguez et al., 2013); the relative positions of the three in silico generated amplimers (black color) encompassing the CpG sites individually interrogated in the present study (pink color); the relative positions of the restriction enzyme sites susceptible to DNA methylation (light blue color ticks); the overall CpG sites across the region (light green ticks); the methylation status at the CpG sites (golden ticks) across a 40-kb long-span view reported in public methylomes from gametes (oocyte and sperm) and somatic tissues (blood, spleen, lung, thymus and liver). The methylation levels are represented on a scale from 0 to 1 (hypomethylated to hypermethylated). Note the overall hypermethylation statuses of the CpGs spanning the *CTLA4* gene in DNA from oocytes, sperm, and somatic tissues. Screenshot prepared using public and custom hubs available from the UCSC Genome Browser (Kent et al., 2002; Raney et al., 2014). (**B**) Zoom-in screenshot across a 460-bp long-span view (hg19; chr2:204734281-204734720), encompassing the 440bp long amplimer that contains the *CTLA4* CpG-1 site interrogated in the study. The annotated features are as in panel (**A**). (**C**) Zoom-in screenshot across a 460-bp long-span view (hg19; chr2:204738666-204739105), encompassing the 440bp long amplimer that contains the *CTLA4* CpG-2 site interrogated in the study. The annotated features are as in panel (**A**)

**FIGURE S3. The *IL1B* locus and the associated CpG sites interrogated**. (**A**) Screenshot across a 40-kb long-span view (hg19; chr2:113573459-113613491) centered at the *IL1B* gene locus. The annotated features are (from top to bottom): chromosome 2 ideogram; physical positions and the exon-intron organization of the reference and alternative *IL1B* transcript isoforms (dark blue color), which are transcribed in the forward direction from the plus DNA strand; the *IL1B* APPRIS principal isoform ENST00000263341 (brown color) (Rodriguez et al., 2013); the relative positions of the three in silico generated amplimers (black color) encompassing the CpG sites individually interrogated in the present study (pink color); the relative positions of the restriction enzyme sites susceptible to DNA methylation (light blue color ticks); the overall CpG sites across the region (light green ticks); the methylation status at the CpG sites (golden ticks) across a 40-kb long-span view reported in public methylomes from gametes (oocyte and sperm) and somatic tissues (blood, spleen, lung, thymus and liver). The methylation levels are represented on a scale from 0 to 1 (hypomethylated to hypermethylated). Note the overall hypermethylation statuses of the CpGs spanning the *IL1B* gene in DNA from oocytes, sperm, and somatic tissues. Screenshot prepared using public and custom hubs available from the UCSC Genome Browser (Kent et al., 2002; Raney et al., 2014). (**B**) Zoom-in screenshot across a 280-bp long-span view (hg19; chr2:113594442-113594701), encompassing the 260bp long amplimer that contains the *IL1B* CpG-1 site interrogated in the study. The annotated features are as in panel (**A**)

**FIGURE S4. The *IL6* locus and the associated CpG sites interrogated**. (**A**) Screenshot across a 40-kb long-span view (hg19; chr7:22,749,736-22,789,767) centered at the *IL6* gene locus. The annotated features are (from top to bottom): chromosome 7 ideogram; physical positions and the exon-intron organization of the reference and alternative *IL6* transcript isoforms (dark blue color), which are transcribed in the forward direction from the plus DNA strand; the *IL6* APPRIS principal isoforms ENST00000404625 and ENST00000258743 (brown color) (Rodriguez et al., 2013); the relative positions of the three in silico generated amplimers (black color) encompassing the CpG sites individually interrogated in the present study (pink color); the relative position of the *cis*-eQTL SNP rs35081782 (black color); the relative positions of the restriction enzyme sites susceptible to DNA methylation (light blue color ticks); the overall CpG sites across the region (light green ticks); the methylation status at the CpG sites (golden ticks) across a 40-kb long-span view reported in public methylomes from gametes (oocyte and sperm) and somatic tissues (blood, spleen, lung, thymus and liver). The methylation levels are represented on a scale from 0 to 1 (hypomethylated to hypermethylated). Note the overall hypermethylation statuses of the CpGs spanning the *IL6* gene in DNA from oocytes, sperm, and somatic tissues. Screenshot prepared using public and custom hubs available from the UCSC Genome Browser (Kent et al., 2002; Raney et al., 2014). (**B**) Zoom-in screenshot across a 280-bp long-span view (hg19; chr7:22765187-22765446), encompassing the 260bp long amplimer that contains the *IL6* CpG-1 site interrogated in the study. The annotated features are as in panel (**A**)

**FIGURE S5. The *IL2* locus and the associated CpG sites interrogated**. (**A**) Screenshot across a 40-kb long-span view (hg19; chr4:123356528-123396559) centered at the *IL2* gene locus. The annotated features are (from top to bottom): chromosome 4 ideogram; physical positions and the exon-intron organization of the reference and alternative *IL2* transcript isoforms (dark blue color), which are transcribed in the forward direction from the plus DNA strand; the *IL2* APPRIS principal isoform ENST00000226730 (brown color) (Rodriguez et al., 2013); the relative positions of the three in silico generated amplimers (black color) encompassing the CpG sites individually interrogated in the present study (pink color); the relative positions of the restriction enzyme sites susceptible to DNA methylation (light blue color ticks); the overall CpG sites across the region (light green ticks); the methylation status at the CpG sites (golden ticks) across a 40-kb long-span view reported in public methylomes from gametes (oocyte and sperm) and somatic tissues (blood, spleen, lung, thymus and liver). The methylation levels are represented on a scale from 0 to 1 (hypomethylated to hypermethylated). Note the overall hypermethylation statuses of the CpGs spanning the *IL2* gene in DNA from oocytes, sperm, and somatic tissues. Screenshot prepared using public and custom hubs available from the UCSC Genome Browser (Kent et al., 2002; Raney et al., 2014). (**B**) Zoom-in screenshot across a 280-bp long-span view (hg19; chr4:123374736-123375025), encompassing the 260bp long amplimer that contains the *IL2* CpG-1 site interrogated in the study. The annotated features are as in panel (**A**)

**FIGURE S6. The *IL10* locus and the associated CpG sites interrogated**. (**A**) Screenshot across a 40-kb long-span view (hg19; chr1:206,923,945-206,963,979) centered at the *IL10* gene locus. The annotated features are (from top to bottom): chromosome 1 ideogram; physical positions and the exon-intron organization of the reference and alternative *IL10* transcript isoforms (dark blue color), which are transcribed in the forward direction from the plus DNA strand; the *IL10* APPRIS principal isoform ENST00000423557 (brown color) (Rodriguez et al., 2013); the relative positions of the three in silico generated amplimers (black color) encompassing the CpG sites individually interrogated in the present study (pink color); the relative positions of the restriction enzyme sites susceptible to DNA methylation (light blue color ticks); the overall CpG sites across the region (light green ticks); the methylation status at the CpG sites (golden ticks) across a 40-kb long-span view reported in public methylomes from gametes (oocyte and sperm) and somatic tissues (blood, spleen, lung, thymus and liver). The methylation levels are represented on a scale from 0 to 1 (hypomethylated to hypermethylated). Note the overall hypomethylation statuses of the CpGs spanning the *IL10* gene in DNA from oocytes in contrast to the hypermethylated statuses in sperm and somatic tissues. Therefore, in the *IL10* gene region, there is asymmetrical methylation in gametes (hypomethylation in oocytes and hypermethylation in spermatozoa). Screenshot prepared using public and custom hubs available from the UCSC Genome Browser (Kent et al., 2002; Raney et al., 2014). (**B**) Zoom-in screenshot across a 460-bp long-span view (hg19; chr1:206945771-206946210), encompassing the 440bp long amplimer that contains the *IL10* CpG-1 site interrogated in the study. The annotated features are as in panel (**A**)

**FIGURE S7. The *CST7* locus and the associated CpG sites interrogated**. (**A**) Screenshot across a 40-kb long-span view (hg19; chr20:24916828-24956861) centered at the *CST7* gene locus. The annotated features are (from top to bottom): chromosome 20 ideogram; physical positions and the exon-intron organization of the reference and alternative *CST7* transcript isoforms (dark blue color), which are transcribed in the forward direction from the plus DNA strand; the *CST7* APPRIS principal isoform ENST00000480798 (brown color) (Rodriguez et al., 2013); the relative positions of the three in silico generated amplimers (black color) encompassing the CpG sites individually interrogated in the present study (pink color); the relative positions of the restriction enzyme sites susceptible to DNA methylation (light blue color ticks); the overall CpG sites across the region (light green ticks); the methylation status at the CpG sites (golden ticks) across a 40-kb long-span view reported in public methylomes from gametes (oocyte and sperm) and somatic tissues (blood, spleen, lung, thymus and liver). The methylation levels are represented on a scale from 0 to 1 (hypomethylated to hypermethylated). Note the overall hypomethylation statuses of the CpGs spanning the *CST7* gene in DNA from oocytes in contrast to the hypermethylated statuses in sperm and somatic tissues. Therefore, in the *CST7* gene region, there is asymmetrical methylation in gametes (hypomethylation in oocytes and hypermethylation in spermatozoa). Screenshot prepared using public and custom hubs available from the UCSC Genome Browser (Kent et al., 2002; Raney et al., 2014). (**B**) Zoom-in screenshot across a 460-bp long-span view (hg19; chr20:24931286-24931725), encompassing the 440bp long amplimer that contains the *CST7* CpG-1 site interrogated in the study. The annotated features are as in panel (**A**)

**FIGURE S8. The *TNF* locus and the associated CpG sites interrogated**. (**A**) Screenshot across a 40-kb long-span view (hg19; chr6:31524812-31564844) centered at the *TNF* gene locus. The annotated features are (from top to bottom): chromosome 6 ideogram; physical positions and the exon-intron organization of the reference and alternative *TNF* transcript isoforms (dark blue color), which are transcribed in the forward direction from the plus DNA strand; the *TNF* APPRIS principal isoform ENST00000449264 (brown color) (Rodriguez et al., 2013); the relative positions of the three in silico generated amplimers (black color) encompassing the CpG sites individually interrogated in the present study (pink color); the relative positions of the restriction enzyme sites susceptible to DNA methylation (light blue color ticks); the overall CpG sites across the region (light green ticks); the methylation status at the CpG sites (golden ticks) across a 40-kb long-span view reported in public methylomes from gametes (oocyte and sperm) and somatic tissues (blood, spleen, lung, thymus and liver). The methylation levels are represented on a scale from 0 to 1 (hypomethylated to hypermethylated). Note the overall hypomethylation statuses of the CpGs spanning the *TNF* gene in DNA from oocytes in contrast to the hypermethylated statuses in sperm and somatic tissues. Therefore, in the *TNF* gene region, there is asymmetrical methylation in gametes (hypomethylation in oocytes and hypermethylation in spermatozoa). Screenshot prepared using public and custom hubs available from the UCSC Genome Browser (Kent et al., 2002; Raney et al., 2014). (**B**) Zoom-in screenshot across a 280-bp long-span view (hg19; chr6:31543454-31543713), encompassing the 260bp long amplimer that contains the *TNF* CpG-1 site interrogated in the study. The annotated features are as in panel (**A**)

**FIGURE S9. The *IFNG* locus and the associated CpG sites interrogated**. (**A**) Screenshot across a 40-kb long-span view (hg19; chr12:68533121-68573153) centered at the *IFNG* gene locus. The annotated features are (from top to bottom): chromosome 12 ideogram; physical positions and the exon-intron organization of the reference and alternative *IFNG* transcript isoforms (dark blue color), which are transcribed in the forward direction from the plus DNA strand; the *IFNG* APPRIS principal isoform ENST00000229135 (brown color) (Rodriguez et al., 2013); the relative positions of the three in silico generated amplimers (black color) encompassing the CpG sites individually interrogated in the present study (pink color); the relative positions of the restriction enzyme sites susceptible to DNA methylation (light blue color ticks); the overall CpG sites across the region (light green ticks); the methylation status at the CpG sites (golden ticks) across a 40-kb long-span view reported in public methylomes from gametes (oocyte and sperm) and somatic tissues (blood, spleen, lung, thymus and liver). The methylation levels are represented on a scale from 0 to 1 (hypomethylated to hypermethylated). Note the overall hypomethylation statuses of the CpGs spanning the *IFNG* gene in DNA from oocytes in contrast to the hypermethylated statuses in sperm and somatic tissues. Therefore, in the *IFNG* gene region, there is asymmetrical methylation in gametes (hypomethylation in oocytes and hypermethylation in spermatozoa). Screenshot prepared using public and custom hubs available from the UCSC Genome Browser (Kent et al., 2002; Raney et al., 2014). (**B**) Zoom-in screenshot across a 460-bp long-span view (hg19; chr12:68553410-68553669), encompassing the 440bp long amplimer that contains the *IFNG* CpG-1 site interrogated in the study. The annotated features are as in panel (**A**)

**FIGURE S10. Representative MSRE-PCR triplex assays for determination of the methylation statuses at the target CpG sites**. Shown are the electrophoretic profiles of the amplimers before and after digestion with the methylation-sensitive *HpyCH4IV* restriction enzyme. (**A**) Hypermethylated status at the *IL4* (CpG-3) site. (**B**) Intermediate methylation at the *IL4*(CpG-1) site. (**C**) Hypomethylated status at the *IL2* (CpG-1) site. In each triplet amplimer assay, the profiles generated from undigested DNA consists of two control products (left and right peaks) and a test product containing the target CpG site (peak in the middle). The control peaks represent the amplimers of a chromosomal region known to be 100% unmethylated in the human genome (left peak) (see Materials and Methods for details) and a chromosomal region lacking *HpyCH4IV* sites (right peak). (**D**) Intermediate methylation status of the rs35081782 insertion/deletion variant alleles in the *IL6* test amplimer from a heterozygote, exhibiting two peaks representing the two variant alleles, one with and the other without the TC insertion. Each panel corresponds to a DNA sample from a different hemophilia A patient.

**FIGURE S11. The range of the methylation levels at the thirteen CpG sites in the discovery subset of subjects**. The methylation statuses are classified in three possible patterns: hypermethylated, intermediate methylated and hypomethylated. The three different groups of subjects are shown on the left. Names of the individual CpG sites are displayed on the right. Each CpG site is depicted in a different color.

**FIGURE S12**. Distribution of the methylation statuses at known blood cell-type specific CpG sites in public methylomes. The x-axis represents the extent of methylation at the individual CpG sites shown on the right. The methylome studies (see Table S4 Dataset (**B**) for details) are named on the left according to the blood-derived cell type, whole blood or gametes.

**Table S1**. Information about the hemophilia A patients included in the study

**Table S2**. Information about the healthy non-hemophilia A subjects included in the study

**Table S3**. Primer sequences, estimated amplimers, physical coordinates of CpG sites and methylation resistant enzymes used in the study

**Table S4**. Dataset (**A**): Details of public methylome studies used for computational cross-referencing. Dataset (**B**): Cross-referenced methylation levels at the individual selected CpG sites interrogated in this study and at CpG sites known to be differentially methylated in blood cell types

**Table S5**. The significance of associations between inhibitor development and levels of methylation at five individual CpG sites

**Table S6**. Genotypes of the *IL6* rs35081782 indel variant in the studied groups of subjects

**Table S7**. Genome-wide significance of associations between the *IL4* rs2227282, *IL6* rs35081782 and human diseases and traits from cross-referencing with genotype-phenotype genome-wide associations studies (GWAS) from PhenoScanner (Dataset **A**); HaploReg lookup of regulatory chromatin states at the lead variant IL4 rs2227282 (Dataset **B**); associated traits from HaploReg (Dataset **C**), and predicted network of protein-protein interactions from STRING database (Dataset **D**)

