## Supplementary Tables and Figures for "A Differentially Methylated CpG Site in the *IL4* Gene Associated with Anti-FVIII Inhibitor Antibody Development in Hemophilia A"

**Table S1. Characterization of the hemophilia A patients included in the study**

| Identification code | Age (years) at collection time | Anti-FVIII inhibitor antibody status | Hemophilia A causative mutation | Mean age (years) | Median age (years) |
|---|---|---|---|---|---|
| 2 | 24 | - | Inv22-1 | | |
| 66 | 21 | - | Inv22-1 | | |
| 123 | 25 | - | Inv22-1 | | |
| 131 | 30 | - | Inv22-1 | | |
| 266 | 12 | - | Inv22-1 | | |
| 298 | 10 | - | Inv22-1 | | |
| 537 | 12 | - | Inv22-2 | | |
| 737 | 15 | - | Inv22-1 | | |
| 756 | 8 | - | Inv22-2 | | |
| 757 | 3 | - | Inv22-2 | | |
| 764 | 2 | - | Inv22-1 | | |
| 826 | 16 | - | Inv22-1 | | |
| 893 | 1 | - | Inv22-1 | | |
| 913 | 5 | - | Inv22-1 | | |
| 1017 | 2 | - | Inv22-1 | | |
| 1048 | 18 | - | Inv22-1 | | |
| 1080 | 2 | - | Inv22-1 | | |
| 1085 | 1 | - | Inv22-1 | | |
| 1145 | 51 | - | Inv22-1 | | |
| 65 | 19 | - transient | Inv22-1 | 13.85 | 12 |
| 1 | 19 | +(HR) | Inv22-1 | | |
| 125 | 27 | +(HR) | Inv22-1 | | |
| 466 | 20 | +(HR) | Inv22-1 | | |
| 501 | 10 | +(HR) | Inv22-1 | | |
| 508 | 32 | +(HR) | Inv22-1 | | |
| 522 | 24 | +(HR) | Inv22-1 | | |
| 600 | 7 | +(HR) | Inv22-1 | | |
| 610 | 5 | +(HR) | Inv22-1 | | |
| 642 | 5 | +(HR) | Inv22-1 | | |
| 654 | 3 | +(HR) | Inv22-1 | | |
| 666 | 10 | +(HR) | Inv22-1 | | |
| 958 | 2 | +(HR) | Inv22-2 | | |
| 961 | 10 | +(HR) | Inv22-1 | | |
| 1055 | 2 | +(HR) | Inv22-2 | | |
| 1109 | 4 | +(HR) | Inv22-1 | | |
| 773 | 1 | +(LR) | Inv22-1 | | |
| 854 | 2 | +(HR) | Inv22-1 | | |
| 879 | 17 | +(HR) | Inv22-1 | | |
| 885 | 46 | +(HR) | Inv22-1 | | |
| 985 | 2 | +(HR) | Inv22-2 | 12.4 | 8.5 |
| | | **Mean age (n=40)** | 13.125 | | |
| | | **Median age (n=40)** | 10 | | |

**Notes:**

+ (LR): hemophilic patient low responder inhibitor

- : hemophilic patient inhibitor negative

+ (HR): hemophilic patient high responder inhibitor

**Table S2. Characterization of healthy non-hemophilia A subjects include in the study**

| Identification Code | Collection date | Age (years) at collection time |
|---|---|---|
| CN30 | 05/23/2011 | 49 |
| CN33 | 05/23/2011 | 42 |
| CN4 | 09/29/2010 | 42 |
| CNv107 | 05/26/2011 | 32 |
| CNv110 | 05/26/2011 | 79 |
| CNv125 | 05/26/2011 | 39 |
| CNv138 | 05/26/2011 | 26 |
| CNv143 | 05/26/2011 | 59 |
| CNv148 | 05/26/2011 | 57 |
| CNv154 | 05/26/2011 | 69 |
| CNv161 | 05/26/2011 | 68 |
| CNv163 | 05/26/2011 | 7 |
| CNv180 | 05/26/2011 | 39 |
| CNv181 | 05/26/2011 | 79 |
| CNv184 | 05/26/2011 | 65 |
| CNv186 | 05/26/2011 | 58 |
| CNv213 | 05/26/2011 | 45 |
| CNv236 | 05/26/2011 | 54 |
| CNv242 | 05/26/2011 | 78 |
| CNv44 | 05/26/2011 | 59 |
| | **Mean age (years)** | **52.3** |
| | **Median age (years)** | **55.5** |

**Table S3. Primer sequences, estimated amplicons, physical coordinates of CpG sites and methylation resistant enzymes used in the study**

| CpG site gene target | Forward Primer (5´FAM labelled) | Reverse Primer | Enzyme | Amplimer size (bp) | Amplimer physical coordinate (hg19) | Individual CpG sites physical coordinates (hg19) |
|---|---|---|---|---|---|---|
| Negative control amplimer | GAGGCAGGAGAACACTGGAG | AAGGACTTCTGCCCCCTAAT | no restriction enzyme | 450 | chr5:132,016,949-132,017,398 | |
| Positive control amplimer | CAGCGGCTTCCTCCTAGC | GGAGAACGAGTCAGTGGCTTC | HpaII, HpyCH4IV | 253 | chr8:27,632,030-27,632,282 | |
| *IL4* (CpG-1) | AACTGCTTCCCCCTCTGTTC | CTTGGAGGCAGCAAAGATGT | HpaII, HpyCH4IV | 440 | chr5:132,009,759-132,010,198 | chr5:132,009,947-132,009,948 |
| *IL4* (CpG-2) | GCATGTTTTGGTCTTTCTGG | AAAATGAAAGCTCCCTCAACC | HpyCH4IV | 260 | chr5:132,010,354-132,010,613 | chr5:132,010,490-132,010,491 |
| *IL4* (CpG-3) | TCCATAATGAACCTCAAATACCTC | AAGGGCAGCTTTAGTGCAAG | HpaII, HpyCH4IV | 440 | chr5:132,012,774-132,013,213 | chr5:132,013,185-132,013,186 |
| *IL6* | CAGTGGCTTCGTTTCATGC | GTGACCTCTGTTGGGCATTT | HpyCH4IV | 260 | chr7:22,765,187-22,765,446 | chr7:22,765,236-22,765,237 |
| *IL1B* | TCACAATCAAGTTAAAGGAAAGG | GTCTTCCACTTTGTCCCACA | HpyCH4IV | 260 | chr2:113,594,442-113,594,701 | chr2:113,594,654-113,594,655 |
| *IL2* | ACATGCATGGGTACTTTACAAAT | AGGCCACAGAACTGAAACATC | HpyCH4IV | 260 | chr4:123,374,751-123,375,010 | chr4:123,374,885-123,374,886 |
| *IL10* | AGCTGTGCATGCCTTCTTTT | TCTGGAATGGGCAATTTGT | HpaII, HpyCH4IV | 440 | chr1:206,945,771-206,946,210 | chr1:206,946,187-206,946,188 |
| *CST7* | ACCCATGGCTGGCAGAAG | ACCACCCAGGAAAATGACG | HpaII, HpyCH4IV | 440 | chr20:24,931,286-24,931,725 | chr20:24,931,707-24,931,708 |
| *TNF* | GGCAGGTTCTCTTCCTCTCA | GGCACTCACCTCTTCCCTCT | HpaII, HpyCH4IV | 260 | chr6:31,543,454-31,543,713 | chr6:31,543,545-31,543,546 |

| | | | | | | |
|---|---|---|---|---|---|---|
| *IFNG* | TTAAGCCAAAGAAGTTG AAATCAG | ACACCCAAATGCCACAA AAC | HpaII, HpyCH4IV | 440 | chr12:68,553,410-68,553,669 | chr12:68,553,577-68,553,578 |
| *CTLA4* (CpG-1) | ATCTGTGGTGGTCGTTTT CC | ATTTCAGGAAGGCAGAT CAAAA | AciI | 440 | chr2:204,738,666-204,739,105 | chr2:204,739,001-204,739,002 |
| *CTLA4* (CpG-2) | AATTGGATCATGGGGGA CTC | TCCCTGGCATTGTTGTAG AG | AciI | 440 | chr2:204,734,281-204,734,720 | chr2:204,734,588-204,734,589 |
| *CD28* | GCAAAATTGAAGTTATG TATCCTCCT | TCTTCCTGAGTCTTAACC CATTAGA | AciI | 440 | chr2:204,591,638-204,592,077 | chr2:204,591,638-204,592,077 |

*FAM prefix represents blue fluorophore-labeled primer.

**Table S4 Dataset (A). Details of public methylome studies used for computational cross-referencing**

| Data Name | Cell Type | Publication | Data Type | Track Name | BS rate | Methylation | Coverage | %CpGs | #HMR | #AMR | #PMD |
|---|---|---|---|---|---|---|---|---|---|---|---|
| Human Oocyte | Mature oocyte cells | Okae et al, 2014 | methylation level | Human_Oocyte_Meth | 1.000 | 0.900 | 10.000 | 1.000 | 0 | 0 | 0 |
| Human Sperm | Mature sperm cells from two donors | Molaro-Sperm-2011 | methylation level | Sperm Methylation Profiles of Human and Chimp, Molaro 2011 : Human_Sperm_Meth | 1.000 | 0.709 | 17.033 | 0.960 | 84906 | 9363 | 0 |
| Human CD4T 100yr | CD4+ T cells (100 year old) | Heyn-Human-NewbornCentenarian-2012 | methylation level | Distinct Human DNA Methylomes from Different Ages, Heyn 2012 : Human_CD4T-100yr_Meth | 0.995 | 0.710 | 14.042 | 0.953 | 45804 | 2891 | 0 |
| Human CD4T Newborn | CD4+ T cells (newborn) | Heyn-Human-NewbornCentenarian-2012 | methylation level | Distinct Human DNA Methylomes from Different Ages, Heyn 2012 : Human_CD4T-Newborn_Meth | 0.995 | 0.802 | 14.417 | 0.954 | 60165 | 1668 | 0 |

| | | | | | | | | | | | |
|---|---|---|---|---|---|---|---|---|---|---|---|
| Human BCell | BCell: normal | Hodges-Human-2011 | methylation level | Changes in Human Hematopoietic Stem Cells, Hodges 2011 : Human_BCell_Meth | 0.992 | 0.749 | 11.855 | 0.957 | 54694 | 2481 | 0 |
| Human Neut | Neutrophil: normal | Hodges-Human-2011 | methylation level | Changes in Human Hematopoietic Stem Cells, Hodges 2011 : Human_Neut_Meth | 0.992 | 0.752 | 11.602 | 0.958 | 73121 | 2111 | 0 |
| Human BloodHealthy | child healthy blood | Gao-Human-2015 | methylation level | Human_BloodHealthy_Meth | 0.990 | 0.811 | 190.490 | 0.987 | 70170 | 0 | 4635 |
| Human Macrophage | Human CD14 macrophage | Roadmap-Human-2015 | methylation level | Roadmap 2015 : Human_Macrophage_Meth | 0.996 | 0.730 | 36.130 | 0.956 | 77058 | 0 | 0 |
| Human NK | Human CD56 NK cells | Roadmap-Human-2015 | methylation level | Roadmap 2015 : Human_NK_Meth | 0.997 | 0.747 | 26.741 | 0.954 | 56897 | 0 | 0 |
| Human Tcell | Human CD3 T cells | Roadmap-Human-2015 | methylation level | Roadmap 2015 : Human_Tcell_Meth | 0.995 | 0.709 | 34.106 | 0.956 | 51640 | 0 | 0 |
| Human Spleen | Spleen primary tissue | Roadmap-Human-2015 | methylation level | Human_Spleen_Meth | 0.996 | 0.804 | 99.441 | 0.967 | 62765 | 20681 | 0 |

| | | | | | | | | | | | |
|---|---|---|---|---|---|---|---|---|---|---|---|
| Human Liver | Liver | Roadmap-Human-2015 | methylation level | Human_Liver_Meth | 0.995 | 0.792 | 49.478 | 0.958 | 58652 | 0 | 0 |
| Human Lung | Lung | Roadmap-Human-2015 | methylation level | Human_Lung_Meth | 0.996 | 0.809 | 78.184 | 0.960 | 57696 | 0 | 0 |
| Human Thymus | Thymus | Roadmap-Human-2015 | methylation level | Human_Thymus_Meth | 0.995 | 0.780 | 129.039 | 0.972 | 59041 | 0 | 0 |
| Human_CD8T cell | Primary CD8 T cell | Komori-Human-2015 | methylation level | Human_CD8T cell_Meth | 0.965 | 0.080 | 0.328 | 0.010 | 2414 | 0 | 0 |

**References**

Roadmap Epigenomics, C., Kundaje, A., Meuleman, W., Ernst, J., Bilenky, M., Yen, A., Heravi-Moussavi, A., Kheradpour, P., Zhang, Z., Wang, J., Ziller, M.J., Amin, V., Whitaker, J.W.,

**Table S4 Dataset (B). Cross-referenced methylation levels at the individual selected CpG sites interrogated in this study and at CpG sites known to be differentially methylated in blood cell types**

| | Gene | CpG site | Chr | Start position | End position | Oocyte | Sperm | Blood Healthy | Liver | Lung | Thymus | Spleen | CD4T 100yr | CD4T Newbo | Neutrophils | B-cell | Macrophages | NK-cell | T-cell | CD8T-cell |
|---|---|---|---|---|---|---|---|---|---|---|---|---|---|---|---|---|---|---|---|---|
| This study | *IL10* | CpG-1 | chr1 | 2.1E+08 | 2.1E+08 | 0 | 0.8636 | 0.3318 | 0.9032 | 0.6238 | 0.2976 | 0.4492 | 0.88 | 0.7037 | 0 | 0.2308 | 0 | 0.48 | 0.875 | 1 |
| | *IFNG* | CpG-1 | chr12 | 6.9E+07 | 6.9E+07 | 0 | 0.8421 | 0.7492 | 0.6441 | 0.7442 | 0.9195 | 0.7514 | 0.5417 | 0.9474 | 0.8462 | 0.9091 | 0.8438 | 0.4 | 0.4783 | 0 |
| | *IL1B* | CpG-1 | chr2 | 1.1E+08 | 1.1E+08 | 1 | 0.7 | 0.1365 | 0.8116 | 0.427 | 0.4828 | 0.2826 | 0.4 | 0.6364 | 0 | 0.2353 | 0 | 0.25 | 0.5 | 0 |
| | *CD28* | CpG-1 | chr2 | 2E+08 | 2E+08 | 0 | 0.4211 | 0.957 | 0.9714 | 1 | 0.9677 | 0.9724 | 0.9231 | 0.9091 | 0.8947 | 0.7931 | 1 | 1 | 1 | 0 |
| | *CD28* | CpG-1 | chr2 | 2E+08 | 2E+08 | 0 | 0.1765 | 0.985 | 0.9722 | 0.989 | 1 | 0.9858 | 1 | 0.9545 | 0.8462 | 0.9615 | 1 | 1 | 1 | 0 |
| | *CTLA4* | CpG-1 | chr2 | 2E+08 | 2E+08 | 1 | 0.85 | 0.8554 | 0.8772 | 0.8983 | 0.8692 | 0.8881 | 0.8462 | 0.6364 | 0.9474 | 0.75 | 0.9818 | 0.6053 | 0.597 | 0 |
| | *CTLA4* | CpG-2 | chr2 | 2E+08 | 2E+08 | 1 | 0.875 | 0.5362 | 0.6 | 0.5088 | 0.525 | 0.5072 | 0.1176 | 0.0526 | 0.5 | 0.8889 | 0.6429 | 0.0606 | 0.0213 | 0 |
| | *CST7* | CpG-1 | chr20 | 2.5E+07 | 2.5E+07 | 0 | 0.9259 | 0.4456 | 0.775 | 0.5652 | 0.8933 | 0.5679 | 0.8235 | 0.8824 | 0.1765 | 0.6364 | 0.6032 | 0.42 | 0.7414 | 0 |
| | *IL2* | CpG-1 | chr4 | 1.2E+08 | 1.2E+08 | 1 | 1 | 0.0474 | 0.494 | 0.1885 | 0.0225 | 0.146 | 0 | 0.0667 | 0 | 0.0526 | 0 | 0 | 0 | 0 |
| | *IL4* | CpG-1 | chr5 | 1.3E+08 | 1.3E+08 | 0 | 0.9032 | 0.5826 | 0.62 | 0.6067 | 0.3038 | 0.5182 | 0.3571 | 0.4167 | 0.6471 | 0.6667 | 0.8378 | 0.5625 | 0.1795 | 0 |
| | *IL4* | CpG-2 | chr5 | 1.3E+08 | 1.3E+08 | 0 | 0.7857 | 0.7261 | 0.3871 | 0.557 | 0.75 | 0.5492 | 0.4091 | 0.9333 | 0.8571 | 0.5714 | 0.7059 | 0.7647 | 0.6923 | 0 |
| | *IL4* | CpG-3 | chr5 | 1.3E+08 | 1.3E+08 | 0 | 0.8696 | 0.9333 | 0.7571 | 0.6458 | 0.907 | 0.7266 | 0.9231 | 0.9474 | 1 | 0.9545 | 0.8793 | 0.9545 | 0.8868 | 0 |
| | *TNF* | CpG-1 | chr6 | 3.2E+07 | 3.2E+07 | 0.2362 | 0.7778 | 0.0614 | 0.2917 | 0.3333 | 0.2037 | 0.125 | 0.1111 | 0.1429 | 0 | 0 | 0 | 0.0645 | 0.0635 | 0 |
| | *IL6* | CpG-1 | chr7 | 2.3E+07 | 2.3E+07 | 1 | 0 | 0.4327 | 0.5763 | 0.7143 | 0.8095 | 0.358 | 0.25 | 1 | 0.5 | 0.5714 | 0.4 | 0.4615 | 0.4333 | 0 |
| cell types[1] | *intergenic* | cg27582527 | chr1 | 1.6E+07 | 1.6E+07 | 0 | 0.8333 | 0.7168 | 0.8846 | 0.7907 | 0.9362 | 0.8512 | 0.75 | 0.8696 | 0.8947 | 0.75 | 0.9265 | 0.2963 | 0.6508 | 0 |
| | *CD8A/RMND* | cg25939861 | chr2 | 8.7E+07 | 8.7E+07 | 0 | 0.9286 | 0.8173 | 0.9344 | 0.9592 | 0.2637 | 0.9055 | 0.84 | 0.9583 | 0.8824 | 0.9444 | 0.9375 | 0.7179 | 0.6098 | 0 |

| | | | | | | | | | | | | | | | | | | | | |
|---|---|---|---|---|---|---|---|---|---|---|---|---|---|---|---|---|---|---|---|---|
| Blood-derived c | *PARK2* | cg23244761 | chr6 | 1.6E+08 | 1.6E+08 | 0.8268 | 0.7297 | 0.8452 | 0.6 | 0.2623 | 0.8725 | 0.6204 | 0.8947 | 0.9286 | 0.8889 | 0.9091 | 0.037 | 0.9474 | 0.9265 | 0 |
| | *intergenic* | cg19276014 | chr10 | 1.3E+08 | 1.3E+08 | 1 | 0.8571 | 0.9021 | 0.9667 | 0.954 | 0.9222 | 0.8081 | 1 | 1 | 0.9583 | 0 | 0.9111 | 0.7143 | 0.8889 | 0 |
| | *WDR20* | cg05398700 | chr14 | 1E+08 | 1E+08 | 1 | 0.6818 | 0.4715 | 0.9512 | 0.8281 | 0.9767 | 0.7353 | 0.75 | 0.7692 | 0 | 1 | 0.8571 | 1 | 1 | 0 |

**[1] References**

Houseman, E.A., Accomando, W.P., Koestler, D.C., Christensen, B.C., Marsit, C.J., Nelson, H.H., et al. (2012). DNA methylation arrays as surrogate measures of cell mixture distribution.BMC Bioinformatics 13, 86. doi: 10.1186/1471-2105-13-86.

**Table S5. Significance of associations between inhibitor development and levels of methylation at five individual CpG sites**

| | Inhibitor (+) versus Inhibitor (-) | | Inhibitor (+) versus Non-hemophilic | | Inhibitor (-) versus Non-hemophilic | | |
|---|---|---|---|---|---|---|---|
| **Individual CpG sites** | ***p-value*** | **95% CI** | ***p-value*** | **95% CI** | ***p-value*** | **95% CI** | **Normal distribution** |
| *IL4* (CpG-1) | 0.802 | -0.07692702 : 0.09874702 | 0.67 | -0.09504200 : 0.06191621 | 0.458 | -0.04694861 : 0.10189440 | YES |
| *IL4* (CpG-2) | 0.241 | -1.857136e-05 : 1.266307e-02 | 0.207 | -3.744918e-05 : 2.137548e-02 | 0.818 | -0.003057561 : 0.021252121 | NO |
| *IL4* (CpG-3) | 0.044 | -1.009419e-01 : 5.148617e-05 | 0.232 | -5.456763e-02 : 1.256362e-05 | 0.6 | -4.30643e-05 : 1.03274e-03 | NO |
| *IFNG* | 0.061 | -1.941792e-01 : 2.372708e-05 | 0.283 | -1.431504e-01 : 6.726593e-05 | 0.654 | -3.347093e-05 : 9.427221e-02 | NO |
| *IL6* | 0.903 | -0.1100508 : 0.1249220 | 0.379 | -0.13731331 : 0.06143067 | 0.507 | -0.15375540 : 0.06498443 | YES |
| *IL2* | 0.212 | -0.05925674 : 0.01819320 | 0.625 | -0.04730356 : 0.01752929 | 0.563 | -0.02923711 : 0.05193650 | NO |
| *IL1B* | 0.427 | -0.08563773 : 0.04516611 | 0.755 | -0.08461738 : 0.05602001 | 0.755 | -0.05654652 : 0.10190133 | NO |
| *IL10* | 0.505 | -0.1314061 : 0.2200554 | 0.733 | -0.2331608 : 0.1104471 | 0.306 | -0.33812657 : 0.08680369 | NO |
| *CST7* | 0.631 | -0.09799805 : 0.15957384 | 0.27 | -0.05468701 : 0.18981701 | 0.54 | -0.15732218 : 0.08376797 | YES |
| *TNF* | 0.85 | -0.05545338 : 0.07165488 | 1 | -0.06137639 : 0.05294737 | 0.733 | -0.06714464 : 0.04778082 | NO |
| *CTLA4* (CpG-1) | 0.677 | -0.12568087 : 0.07407054 | 0.384 | -0.11422237 : 0.05149954 | 0.909 | -0.10576336 : 0.06897295 | YES |
| *CTLA4* (CpG-2) | 0.5 | -0.1462564 : 0.1163737 | 0.122 | -1.558348e-01 : 4.869991e-05 | 0.638 | -0.16112623 : 0.01702729 | NO |
| *CD28* | 0.376 | -6.603508e-05 : 7.658077e-02 | 0.608 | -7.677285e-05 : 1.102532e-02 | 0.701 | -0.05833449 : 0.01096423 | NO |

| ***IL4* CpG-3** | **Mean** | **Standard Deviation** |
|---|---|---|
| Inhibitor positive | 0.9153125 | 0.09478952 |
| Inhibitor negative | 0.9510889 | 0.1160409 |
| Non-hemophilic | 0.93777 | 0.1038556 |
| ***IFNG* CpG** | **Mean** | **Standard Deviation** |
| Inhibitor positive | 0.8939125 | 0.1490099 |
| Inhibitor negative | 0.97754 | 0.06860049 |
| Non-hemophilic | 0.95828 | 0.0738304 |

**Table S6. Genotypes of the *IL6* rs35081782 indel variant in the studied groups of subjects**

| Hemophilia A patients with anti-FVIII inhibitor antibodies | | | |
|---|---|---|---|
| Minor allele (256) frequency: 15% | | | |
| **Sample code** | ***IL6* Genotype (bp)** | | **Methylation (%)** |
| 885 | 256 | 258 | 41.66 |
| 522 | 258 | 258 | 45.01 |
| 125 | 258 | 258 | 58.9 |
| 1109 | 258 | 258 | 63.81 |
| 600 | 258 | 258 | 63.82 |
| 1 | 258 | 258 | 64.09 |
| 854 | 258 | 258 | 65.42 |
| 508 | 258 | 258 | 66.47 |
| 961 | 256 | 258 | 68.28 |
| 958 | 256 | 258 | 71.62 |
| 642 | 258 | 258 | 76.28 |
| 466 | 256 | 258 | 83.08 |
| 666 | 258 | 258 | 85.36 |
| 654 | 258 | 258 | 85.6 |
| 501 | 258 | 258 | 87.42 |
| 985 | 258 | 258 | 92.67 |
| 879 | 256 | 258 | 93.46 |
| 773 | 256 | 258 | 96.57 |
| 610 | 258 | 258 | 100 |
| 1055 | 258 | 258 | 100 |

| Hemophilia A patients without anti-FVIII inhibitor antibodies | | | |
|---|---|---|---|
| Minor allele (256) frequency: 27.5% | | | |
| **Sample code** | ***IL6* Genotype (bp)** | | **Methylation (%)** |
| 537 | 258 | 258 | 33.71 |
| 66 | 256 | 258 | 51.16 |
| 65 | 256 | 258 | 61.54 |
| 756 | 258 | 258 | 62.18 |
| 1145 | 256 | 258 | 62.22 |
| 123 | 258 | 258 | 64.1 |
| 2 | 258 | 258 | 68.8 |
| 764 | 256 | 258 | 69.36 |
| 1048 | 258 | 258 | 69.9 |
| 737 | 258 | 258 | 71.72 |
| 757 | 256 | 258 | 72.87 |
| 1017 | 256 | 258 | 74.88 |
| 131 | 256 | 258 | 82.62 |
| 298 | 258 | 258 | 84.4 |
| 826 | 258 | 258 | 90.52 |
| 893 | 258 | 258 | 94.93 |
| 266 | 256 | 258 | 100 |
| 913 | 256 | 256 | 100 |
| 1080 | 258 | 258 | 100 |
| 1085 | 256 | 258 | 100 |

| Healthy non-hemophiliac subjects | | | |
|---|---|---|---|
| Minor allele (256) frequency: 25% | | | |
| **Sample code** | ***IL6* Genotype (bp)** | | **Methylation (%)** |
| 143 | 258 | 258 | 52.74 |
| 154 | 256 | 258 | 59.75 |
| 33 | 256 | 258 | 63.99 |
| 236 | 258 | 258 | 67.6 |
| 107 | 258 | 258 | 68.47 |
| 213 | 256 | 258 | 70.32 |

| | | | |
|---|---|---|---|
| 110 | 256 | 258 | 70.33 |
| 30 | 256 | 258 | 73.67 |
| 44 | 258 | 258 | 75.74 |
| 4 | 258 | 258 | 76.13 |
| 184 | 256 | 258 | 77.55 |
| 181 | 258 | 258 | 87.32 |
| 161 | 256 | 258 | 89.36 |
| 163 | 256 | 258 | 90.97 |
| 148 | 258 | 258 | 92.24 |
| 242 | 256 | 258 | 92.61 |
| 186 | 258 | 258 | 96.61 |
| CN 125 | 256 | 258 | 97.26 |
| 180 | 258 | 258 | 97.4 |
| 138 | 258 | 258 | 100 |

| **Chi square test** | **p-value** |
|---|---|
| Inh(+)/Inh(-) | 0.17 |
| Inh(+)/Non-h | 0.26 |
| Inh(-)/Non-h | 0.95 |

Table S7 Dataset (A): PhenoScanner lookup for GWAS

| SNP | Physical coordinates (hg19) | Allele 1 | Allele 2 | Associated trait * | Beta | SE | *p* -value | Direction | sample (n) | Cases (n) | Controls (n) | Studies (n) | Study dataset | Population ancestry | Publication Year | PMID |
|---|---|---|---|---|---|---|---|---|---|---|---|---|---|---|---|---|
| rs2227282 | chr5:132013179 | G | C | Atopic dermatitis | -0.1237 | 0.0195 | 2.3E-10 | - | 1.0E+05 | 1.9E+04 | 8.4E+04 | 22 | EAGLE_Eczema_Mixed_2015 | Mixed | 2015 | 26482879 |
| rs2227282 | chr5:132013179 | G | C | Allergic disease | -0.0368 | 0.0067 | 4.4E-05 | - | 3.6E+05 | 1.8E+05 | 1.8E+05 | 13 | Ferreira-M_Allergic-Disease_EUR_2017 | European | 2017 | 29083406 |
| rs2227282 | chr5:132013179 | G | C | Height | 0.017 | 0.0034 | 5.2E-07 | + | 2.5E+05 | 0.0E+00 | 2.5E+05 | 79 | GIANT_Height_EUR_2014 | European | 2014 | 25282103 |
| rs2227282 | chr5:132013179 | G | C | Asthma | -0.005364 | 0.0009197 | 5.5E-06 | - | 3.4E+05 | 3.9E+04 | 3.0E+05 | 1 | Neale-B_UKBB_EUR_2017 | European | 2017 | UKBB |
| rs2227282 | chr5:132013179 | G | C | No blood clot, bronchitis, emphysema, asthma, rhinitis, eczema or allergy diagnosed by doctor | 0.006658 | 0.001345 | 7.4E-04 | + | 3.4E+05 | 2.3E+05 | 1.1E+05 | 1 | Neale-B_UKBB_EUR_2017 | European | 2017 | UKBB |
| rs2227282 | chr5:132013179 | G | C | Self-reported asthma | -0.005098 | 0.0009214 | 3.2E-05 | - | 3.4E+05 | 3.9E+04 | 3.0E+05 | 1 | Neale-B_UKBB_EUR_2017 | European | 2017 | UKBB |
| rs2227282 | chr5:132013179 | G | C | Self-reported eczema or dermatitis | -0.002392 | 0.0004572 | 1.7E-04 | - | 3.4E+05 | 8.7E+03 | 3.3E+05 | 1 | Neale-B_UKBB_EUR_2017 | European | 2017 | UKBB |

| | | | | | | | | | | | | | | | | |
|---|---|---|---|---|---|---|---|---|---|---|---|---|---|---|---|---|
| rs2227282 | chr5:132013179 | G | C | Time spent using computer | -0.01138 | 0.002541 | 7.5E-03 | - | 2.6E+05 | 0.0E+00 | 2.6E+05 | 1 | Neale-B_UKBB_EUR_2017 | European | 2017 | UKBB |
| rs35081782 | chr7:22765336 | - | TC | Eosinophil count | NA | NA | 8.9E-14 | + | 1.7E+05 | 0.0E+00 | 1.7E+05 | 2 | Astle-W_Blood-Cell-Traits_EUR_2016 | European | 2016 | 27863252 |
| rs35081782 | chr7:22765336 | - | TC | Eosinophil percentage of granulocytes | NA | NA | 1.0E-06 | + | 1.7E+05 | 0.0E+00 | 1.7E+05 | 2 | Astle-W_Blood-Cell-Traits_EUR_2016 | European | 2016 | 27863252 |
| rs35081782 | chr7:22765336 | - | TC | Eosinophil percentage of white cells | NA | NA | 4.3E-09 | + | 1.7E+05 | 0.0E+00 | 1.7E+05 | 2 | Astle-W_Blood-Cell-Traits_EUR_2016 | European | 2016 | 27863252 |
| rs35081782 | chr7:22765336 | - | TC | Granulocyte count | NA | NA | 6.0E-03 | + | 1.7E+05 | 0.0E+00 | 1.7E+05 | 2 | Astle-W_Blood-Cell-Traits_EUR_2016 | European | 2016 | 27863252 |
| rs35081782 | chr7:22765336 | - | TC | Myeloid white cell count | NA | NA | 7.8E-03 | + | 1.7E+05 | 0.0E+00 | 1.7E+05 | 2 | Astle-W_Blood-Cell-Traits_EUR_2016 | European | 2016 | 27863252 |
| rs35081782 | chr7:22765336 | - | TC | Neutrophil percentage of granulocytes | NA | NA | 7.4E-06 | - | 1.7E+05 | 0.0E+00 | 1.7E+05 | 2 | Astle-W_Blood-Cell-Traits_EUR_2016 | European | 2016 | 27863252 |

| | | | | | | | | | | | | | | | | |
|---|---|---|---|---|---|---|---|---|---|---|---|---|---|---|---|---|
| rs35081782 | chr7:22765336 | - | TC | Sum eosinophil basophil counts | NA | NA | 2.4E-13 | + | 1.7E+05 | 0.0E+00 | 1.7E+05 | 2 | Astle-W_Blood-Cell-Traits_EUR_2016 | European | 2016 | 27863252 |
| rs35081782 | chr7:22765336 | - | TC | Sum neutrophil eosinophil counts | NA | NA | 4.4E-06 | + | 1.7E+05 | 0.0E+00 | 1.7E+05 | 2 | Astle-W_Blood-Cell-Traits_EUR_2016 | European | 2016 | 27863252 |
| rs35081782 | chr7:22765336 | - | TC | White blood cell count | NA | NA | 5.4E-03 | + | 1.7E+05 | 0.0E+00 | 1.7E+05 | 2 | Astle-W_Blood-Cell-Traits_EUR_2016 | European | 2016 | 27863252 |

*Source: PhenoScanner. http://www.phenoscanner.medschl.cam.ac.uk/phenoscanner

**Table S7 Dataset (B): HaploReg lookup of the regulatory states at the *IL14* rs2227282 variant**

| #rsID | chr | position (hg19) | Ref/Alt |
|---|---|---|---|
| rs2227282 | chr5 | 132013179 | C/G |

| Epigenome ID (EID) | Group | Description | Chromatin states | Chromatin states | H3K4me1 | H3K4me3 | H3K27ac | H3K9ac | DNase |
|---|---|---|---|---|---|---|---|---|---|
| E017 | IMR90 | IMR90 fetal lung fibroblasts Cell Line | 7_Enh | 15_EnhAF | H3K4me1_Enh | | H3K27ac_Enh | H3K9ac_Pro | |
| E002 | ESC | ES-WA7 Cells | | 19_DNase | | | [redacted] | | [redacted] |
| E008 | ESC | H9 Cells | | 19_DNase | | | H3K27ac_Enh | | |
| E001 | ESC | ES-I3 Cells | 7_Enh | 19_DNase | | | [redacted] | | [redacted] |
| E015 | ESC | HUES6 Cells | | 19_DNase | | | | | [redacted] |
| E014 | ESC | HUES48 Cells | | 19_DNase | | | | | [redacted] |
| E016 | ESC | HUES64 Cells | | 19_DNase | | | | | [redacted] |
| E003 | ESC | H1 Cells | | 19_DNase | | | | | DNase |
| E024 | ESC | ES-UCSF4 Cells | | 19_DNase | | | [redacted] | [redacted] | [redacted] |
| E020 | iPSC | iPS-20b Cells | | 19_DNase | | | | | [redacted] |
| E018 | iPSC | iPS-15b Cells | | 19_DNase | | | [redacted] | H3K9ac_Pro | [redacted] |
| E021 | iPSC | iPS DF 6.9 Cells | | 19_DNase | | | | [redacted] | DNase |
| E022 | iPSC | iPS DF 19.11 Cells | | 19_DNase | | | | [redacted] | |
| E007 | ES-deriv | H1 Derived Neuronal Progenitor Cultured Cells | | 19_DNase | | | | | |
| E009 | ES-deriv | H9 Derived Neuronal Progenitor Cultured Cells | 7_Enh | 19_DNase | H3K4me1_Enh | | [redacted] | [redacted] | [redacted] |
| E010 | ES-deriv | H9 Derived Neuron Cultured Cells | 7_Enh | 19_DNase | H3K4me1_Enh | | [redacted] | [redacted] | [redacted] |

| | | | | | | | | | |
|---|---|---|---|---|---|---|---|---|---|
| E013 | ES-deriv | hESC Derived CD56+ Mesoderm Cultured Cells | 7_Enh | 18_EnhAc | H3K4me1_Enh | | | | |
| E011 | ES-deriv | hESC Derived CD184+ Endoderm Cultured Cells | | 19_DNase | | | | | |
| E004 | ES-deriv | H1 BMP4 Derived Mesendoderm Cultured Cells | | 19_DNase | | | | | |
| E005 | ES-deriv | H1 BMP4 Derived Trophoblast Cultured Cells | | 17_EnhW2 | H3K4me1_Enh | | | | DNase |
| E006 | ES-deriv | H1 Derived Mesenchymal Stem Cells | 7_Enh | 17_EnhW2 | H3K4me1_Enh | | H3K27ac_Enh | | DNase |
| E062 | Blood & T-cell | Primary mononuclear cells from peripheral blood | | 19_DNase | | | | H3K9ac_Pro | |
| E034 | Blood & T-cell | Primary T cells from peripheral blood | | 19_DNase | H3K4me1_Enh | | | | |
| E039 | Blood & T-cell | Primary T helper naive cells from peripheral blood | | 19_DNase | | | | | |
| E041 | Blood & T-cell | Primary T helper cells PMA-I stimulated | | | | H3K4me3_Pro | | | |

| | | | | | | | |
|---|---|---|---|---|---|---|---|
| E042 | Blood & T-cell | Primary T helper 17 cells PMA-I stimulated | | 19_DNase | | | |
| E037 | Blood & T-cell | Primary T helper memory cells from peripheral blood 2 | | 19_DNase | H3K4me1_Enh | | |
| E048 | Blood & T-cell | Primary T CD8+ memory cells from peripheral blood | | 19_DNase | H3K4me1_Enh | | H3K27ac_Enh |
| E038 | Blood & T-cell | Primary T helper naive cells from peripheral blood | | 19_DNase | | | H3K27ac_Enh |
| E050 | HSC & B-cell | Primary hematopoietic stem cells G-CSF-mobilized Female | | 19_DNase | | | |
| E026 | Mesench | Bone Marrow Derived Cultured Mesenchymal Stem Cells | 7_Enh | 15_EnhAF | H3K4me1_Enh | | H3K27ac_Enh |
| E049 | Mesench | Mesenchymal Stem Cell Derived Chondrocyte Cultured Cells | 7_Enh | 17_EnhW2 | H3K4me1_Enh | | |

| | | | | | | | | | |
|---|---|---|---|---|---|---|---|---|---|
| E025 | Mesench | Adipose Derived Mesenchymal Stem Cell Cultured Cells | 7_Enh | 14_EnhA2 | H3K4me1_Enh | H3K4me3_Pro | | | |
| E023 | Mesench | Mesenchymal Stem Cell Derived Adipocyte Cultured Cells | 7_Enh | 17_EnhW2 | H3K4me1_Enh | | | | |
| E052 | Myosat | Muscle Satellite Cultured Cells | 7_Enh | 14_EnhA2 | H3K4me1_Enh | | | H3K9ac_Pro | |
| E055 | Epithelial | Foreskin Fibroblast Primary Cells skin01 | 7_Enh | 14_EnhA2 | H3K4me1_Enh | H3K4me3_Pro | H3K27ac_Enh | | DNase |
| E056 | Epithelial | Foreskin Fibroblast Primary Cells skin02 | | 14_EnhA2 | H3K4me1_Enh | H3K4me3_Pro | H3K27ac_Enh | | |
| E059 | Epithelial | Foreskin Melanocyte Primary Cells skin01 | | 17_EnhW2 | | | | | |
| E061 | Epithelial | Foreskin Melanocyte Primary Cells skin03 | | 17_EnhW2 | | | | | |
| E057 | Epithelial | Foreskin Keratinocyte Primary Cells skin02 | 7_Enh | 16_EnhW1 | H3K4me1_Enh | | | | |
| E058 | Epithelial | Foreskin Keratinocyte Primary Cells skin03 | 7_Enh | 15_EnhAF | H3K4me1_Enh | | | | |

| | | | | | | | |
|---|---|---|---|---|---|---|---|
| E028 | Epithelial | Breast variant Human Mammary Epithelial Cells (vHMEC) | 7_Enh | 16_EnhW1 | H3K4me1_Enh | | |
| E027 | Epithelial | Breast Myoepithelial Primary Cells | | 17_EnhW2 | H3K4me1_Enh | | |
| E054 | Neurosph | Ganglion Eminence derived primary cultured neurospheres | | 19_DNase | | | |
| E053 | Neurosph | Cortex derived primary cultured neurospheres | | 19_DNase | | | |
| E112 | Thymus | Thymus | | 19_DNase | | | |
| E093 | Thymus | Fetal Thymus | 7_Enh | 18_EnhAc | H3K4me1_Enh | H3K4me3_Pro | H3K27ac_Enh |
| E071 | Brain | Brain Hippocampus Middle | | 19_DNase | | | |
| E074 | Brain | Brain Substantia Nigra | | 19_DNase | | | |
| E068 | Brain | Brain Anterior Caudate | | 19_DNase | | | |
| E069 | Brain | Brain Cingulate Gyrus | | 19_DNase | | | |
| E072 | Brain | Brain Inferior Temporal Lobe | | 19_DNase | | | |
| E067 | Brain | Brain Angular Gyrus | | 19_DNase | | | |
| E073 | Brain | Brain_Dorsolateral_Prefrontal_Cortex | | 19_DNase | | | |

| | | | | | | | | |
|---|---|---|---|---|---|---|---|---|
| E082 | Brain | Fetal Brain Female | | 19_DNase | | | | |
| E081 | Brain | Fetal Brain Male | | 19_DNase | | | | |
| E063 | Adipose | Adipose Nuclei | | 17_EnhW2 | H3K4me1_Enh | | | |
| E100 | Muscle | Psoas Muscle | | 19_DNase | | | | |
| E108 | Muscle | Skeletal Muscle Female | | 17_EnhW2 | H3K4me1_Enh | | | |
| E107 | Muscle | Skeletal Muscle Male | | 17_EnhW2 | | | H3K9ac_Pro | |
| E089 | Muscle | Fetal Muscle Trunk | | 19_DNase | H3K4me1_Enh | | | |
| E090 | Muscle | Fetal Muscle Leg | | 19_DNase | H3K4me1_Enh | | | |
| E083 | Heart | Fetal Heart | | 19_DNase | | | | |
| E104 | Heart | Right Atrium | 7_Enh | 19_DNase | H3K4me1_Enh | | | |
| E095 | Heart | Left Ventricle | | 19_DNase | | | | |
| E105 | Heart | Right Ventricle | | 19_DNase | | | | |
| E065 | Heart | Aorta | | 17_EnhW2 | | H3K27ac_Enh | | |
| E078 | Sm. Muscle | Duodenum Smooth Muscle | | 17_EnhW2 | H3K4me1_Enh | H3K27ac_Enh | | |
| E076 | Sm. Muscle | Colon Smooth Muscle | 7_Enh | 16_EnhW1 | H3K4me1_Enh | | H3K9ac_Pro | |
| E103 | Sm. Muscle | Rectal Smooth Muscle | 7_Enh | 17_EnhW2 | H3K4me1_Enh | H3K27ac_Enh | H3K9ac_Pro | |
| E111 | Sm. Muscle | Stomach Smooth Muscle | 7_Enh | 17_EnhW2 | H3K4me1_Enh | | | |
| E092 | Digestive | Fetal Stomach | | 17_EnhW2 | | H3K27ac_Enh | | |
| E085 | Digestive | Fetal Intestine Small | 7_Enh | 17_EnhW2 | H3K4me1_Enh | H3K27ac_Enh | | |
| E084 | Digestive | Fetal Intestine Large | 7_Enh | 17_EnhW2 | H3K4me1_Enh | H3K27ac_Enh | | |
| E109 | Digestive | Small Intestine | | 17_EnhW2 | H3K4me1_Enh | H3K27ac_Enh | | DNase |
| E106 | Digestive | Sigmoid Colon | | 17_EnhW2 | | | | |

| | | | | | | | | | |
|---|---|---|---|---|---|---|---|---|---|
| E075 | Digestive | Colonic Mucosa | | 17_EnhW2 | H3K4me1_Enh | | H3K27ac_Enh | | |
| E101 | Digestive | Rectal Mucosa Donor 29 | | 17_EnhW2 | H3K4me1_Enh | | H3K27ac_Enh | | |
| E102 | Digestive | Rectal Mucosa Donor 31 | | 19_DNase | | | | | |
| E110 | Digestive | Stomach Mucosa | 7_Enh | 17_EnhW2 | H3K4me1_Enh | | | H3K9ac_Pro | |
| E077 | Digestive | Duodenum Mucosa | 7_Enh | 16_EnhW1 | H3K4me1_Enh | | | H3K9ac_Pro | |
| E079 | Digestive | Esophagus | 7_Enh | 17_EnhW2 | H3K4me1_Enh | | H3K27ac_Enh | | |
| E094 | Digestive | Gastric | | 16_EnhW1 | H3K4me1_Enh | H3K4me3_Pro | H3K27ac_Enh | | |
| E099 | Other | Placenta Amnion | | 17_EnhW2 | H3K4me1_Enh | | H3K27ac_Enh | | |
| E086 | Other | Fetal Kidney | | 17_EnhW2 | | | | | |
| E088 | Other | Fetal Lung | | 17_EnhW2 | H3K4me1_Enh | | | | |
| E097 | Other | Ovary | | 17_EnhW2 | | | H3K27ac_Enh | | |
| E087 | Other | Pancreatic Islets | | 19_DNase | | | | | |
| E080 | Other | Fetal Adrenal Gland | 7_Enh | 17_EnhW2 | H3K4me1_Enh | | H3K27ac_Enh | | DNase |
| E091 | Other | Placenta | 7_Enh | 16_EnhW1 | H3K4me1_Enh | | | | DNase |
| E066 | Other | Liver | | 19_DNase | | | | | |
| E098 | Other | Pancreas | | 19_DNase | | | | | |
| E096 | Other | Lung | | 17_EnhW2 | | | H3K27ac_Enh | | |
| E113 | Other | Spleen | | 19_DNase | | | | | |
| E114 | ENCODE, 2012 | A549 EtOH 0.02pct Lung Carcinoma Cell Line | 7_Enh | 14_EnhA2 | H3K4me1_Enh | H3K4me3_Pro | H3K27ac_Enh | H3K9ac_Pro | DNase |
| E115 | ENCODE, 2012 | Dnd41 TCell Leukemia Cell Line | | 17_EnhW2 | | | | | |
| E116 | ENCODE, 2012 | GM12878 Lymphoblastoid Cells | | 19_DNase | | | | | |

| | | | | | | | | | |
|---|---|---|---|---|---|---|---|---|---|
| E117 | ENCODE, 2012 | HeLa-S3 Cervical Carcinoma Cell Line | 7_Enh | 17_EnhW2 | H3K4me1_Enh | | | | DNase |
| E118 | ENCODE, 2012 | HepG2 Hepatocellular Carcinoma Cell Line | | 17_EnhW2 | H3K4me1_Enh | | | | DNase |
| E119 | ENCODE, 2012 | HMEC Mammary Epithelial Primary Cells | 7_Enh | 16_EnhW1 | H3K4me1_Enh | | | | |
| E120 | ENCODE, 2012 | HSMM Skeletal Muscle Myoblasts Cells | 7_Enh | 17_EnhW2 | H3K4me1_Enh | | | | DNase |
| E121 | ENCODE, 2012 | HSMM cell derived Skeletal Muscle Myotubes Cells | 7_Enh | 17_EnhW2 | H3K4me1_Enh | | H3K27ac_Enh | H3K9ac_Pro | DNase |
| E122 | ENCODE, 2012 | HUVEC Umbilical Vein Endothelial Primary Cells | 7_Enh | 17_EnhW2 | H3K4me1_Enh | | | | |
| E123 | ENCODE, 2012 | K562 Leukemia Cells | 7_Enh | 19_DNase | H3K4me1_Enh | | | H3K9ac_Pro | DNase |
| E125 | ENCODE, 2012 | NH-A Astrocytes Primary Cells | 7_Enh | 14_EnhA2 | H3K4me1_Enh | | H3K27ac_Enh | H3K9ac_Pro | DNase |
| E126 | ENCODE, 2012 | NHDF-Ad Adult Dermal Fibroblast Primary Cells | 7_Enh | 17_EnhW2 | H3K4me1_Enh | | | | DNase |

| | | | | | | | | | |
|---|---|---|---|---|---|---|---|---|---|
| E127 | ENCODE, 2012 | NHEK-Epidermal Keratinocyte Primary Cells | 7_Enh | 16_EnhW1 | H3K4me1_Enh | | | | |
| E128 | ENCODE, 2012 | NHLF Lung Fibroblast Primary Cells | 7_Enh | 14_EnhA2 | H3K4me1_Enh | | H3K27ac_Enh | | DNase |
| E129 | ENCODE, 2012 | Osteoblast Primary Cells | 7_Enh | 14_EnhA2 | H3K4me1_Enh | H3K4me3_Pro | H3K27ac_Enh | | |

**missing data**

**Table S7 Dataset (C): HaploReg lookup for GWAS**

| Study ID[1] | Tissue | Correlated gene | *p-value* | Physical coordinates (hg19) |
|---|---|---|---|---|
| Zeller, et al. 2010 | peripheral blood monocytes | *SLC22A4* | 5.55E-09 | chr5:131630145-131679899 |
| Zeller, et al. 2010 | peripheral blood monocytes | *SLC22A5* | 5.35E-07 | chr5:131705401-131731306 |
| GTEx 2015_v6 | Adipose_Subcutaneous | *IL4* | 2.00E-08 | chr5:132009743-132018279 |
| GTEx 2015_v6 | Pituitary | *IL4* | 3.67E-08 | chr5:132009743-132018279 |
| GTEx 2015_v6 | Adipose_Subcutaneous | *KIF3A* | 1.82E-06 | chr5:132028323-132073265 |
| GTEx 2015_v6 | Artery_Tibial | *KIF3A* | 2.35E-05 | chr5:132028323-132073265 |
| GTEx 2015_v6 | Esophagus_Muscularis | *KIF3A* | 8.14E-07 | chr5:132028323-132073265 |
| GTEx 2015_v6 | Nerve_Tibial | *KIF3A* | 2.05E-06 | chr5:132028323-132073265 |
| GTEx 2015_v6 | Skin_Sun_Exposed_Lower_leg | *KIF3A* | 4.68E-07 | chr5:132028323-132073265 |
| GTEx 2015_v6 | Cells_Transformed_fibroblasts | *SEPT8* | 7.20E-08 | chr5:132086509-132113067 |
| Lappalainen, et al. 2013 | Lymphoblastoid_EUR_exonlevel | *PDLIM4-exon 2* | 6.90E-07 | chr5:131598302-131598453 |
| Lappalainen, et al. 2013 | Lymphoblastoid_EUR_exonlevel | *PDLIM4-exon 3* | 6.13E-06 | chr5:131602157-131602238 |
| Lappalainen, et al. 2013 | Lymphoblastoid_EUR_exonlevel | *PDLIM4-exon 4* | 7.75E-08 | chr5:131606608-131606786 |
| Lappalainen, et al. 2013 | Lymphoblastoid_EUR_exonlevel | *PDLIM4-exon 5* | 4.52E-08 | chr5:131606996-131607159 |
| Lappalainen, et al. 2013 | Lymphoblastoid_EUR_exonlevel | *PDLIM4-exon 7* | 4.20E-08 | chr5:131607718-131609147 |
| Lappalainen, et al. 2013 | Lymphoblastoid_EUR_genelevel | *PDLIM4* | 1.17E-07 | chr5:131593351-131609147 |
| Westra, et al. 2013 | Whole_Blood | *IL4* | 6.43E-04 | chr5:132009743-132018279 |
| Westra, et al. 2013 | Whole_Blood | *RAD50* | 0.003357039 | chr5:131892616-131980313 |

[1] **References**

CONSORTIUM, G. T. Human genomics. The Genotype-Tissue Expression (GTEx) pilot analysis: multitissue gene regulation in humans. Science, v. 348, n. 6235, p. 648-660, May 8 2015.

LAPPALAINEN, T. et al. Transcriptome and genome sequencing uncovers functional variation in humans. Nature, v. 501, n. 7468, p. 506-511, Sep 26 2013.

WESTRA, H. J. et al. Systematic identification of trans eQTLs as putative drivers of known disease associations. Nat Genet, v. 45, n. 10, p. 1238-1243, Oct 2013.

ZELLER, T. et al. Genetics and beyond--the transcriptome of human monocytes and disease susceptibility. PLoS One, v. 5, n. 5, p. e10693, May 18 2010.

**Table S7 Dataset (D): Predicted network of protein-protein interactions from STRING database**

| | Biological Process (GO) | | | | |
|---|---|---|---|---|---|
| **Pathway ID** | **Pathway description** | **Count in gene set** | **False discovery rate** | **Matching proteins the network (labels)** | **Matching proteins the network (IDs)** |
| GO:1902603 | carnitine transmembrane transport | 2 | 0.0253 | SLC22A4,SLC22A5 | ENSP00000200652,ENSP00000245407 |
| GO:0015695 | organic cation transport | 2 | 0.029 | SLC22A4,SLC22A5 | ENSP00000200652,ENSP00000245407 |

| | Molecular Function (GO) | | | | |
|---|---|---|---|---|---|
| **Pathway ID** | **Pathway description** | **Count in gene set** | **False discovery rate** | **Matching proteins the network (labels)** | **Matching proteins the network (IDs)** |
| GO:0015226 | carnitine transmembrane transporter activity | 2 | 0.000619 | SLC22A4,SLC22A5 | ENSP00000200652,ENSP00000245407 |
| GO:0032550 | purine ribonucleoside binding | 5 | 0.0239 | KIF3A,RAD50,SEPT8,SLC22A4,SLC22A5 | ENSP00000200652,ENSP00000245407,ENSP00000265335,ENSP00000367991,ENSP00000368020 |
| GO:0032555 | purine ribonucleotide binding | 5 | 0.0239 | KIF3A,RAD50,SEPT8,SLC22A4,SLC22A5 | ENSP00000200652,ENSP00000245407,ENSP00000265335,ENSP00000367991,ENSP00000368020 |
| GO:0035639 | purine ribonucleoside triphosphate binding | 5 | 0.0239 | KIF3A,RAD50,SEPT8,SLC22A4,SLC22A5 | ENSP00000200652,ENSP00000245407,ENSP00000265335,ENSP00000367991,ENSP00000368020 |
| GO:0030165 | PDZ domain binding | 2 | 0.0371 | SLC22A4,SLC22A5 | ENSP00000200652,ENSP00000245407 |

**Note**: Using STRING v11 database, there were no significant pathway enrichments observed in the following categories: Cellular Component (GO), KEGG Pathways, PFAM Protein

**Predicted network of interactions (low score confidence = 0.3)**

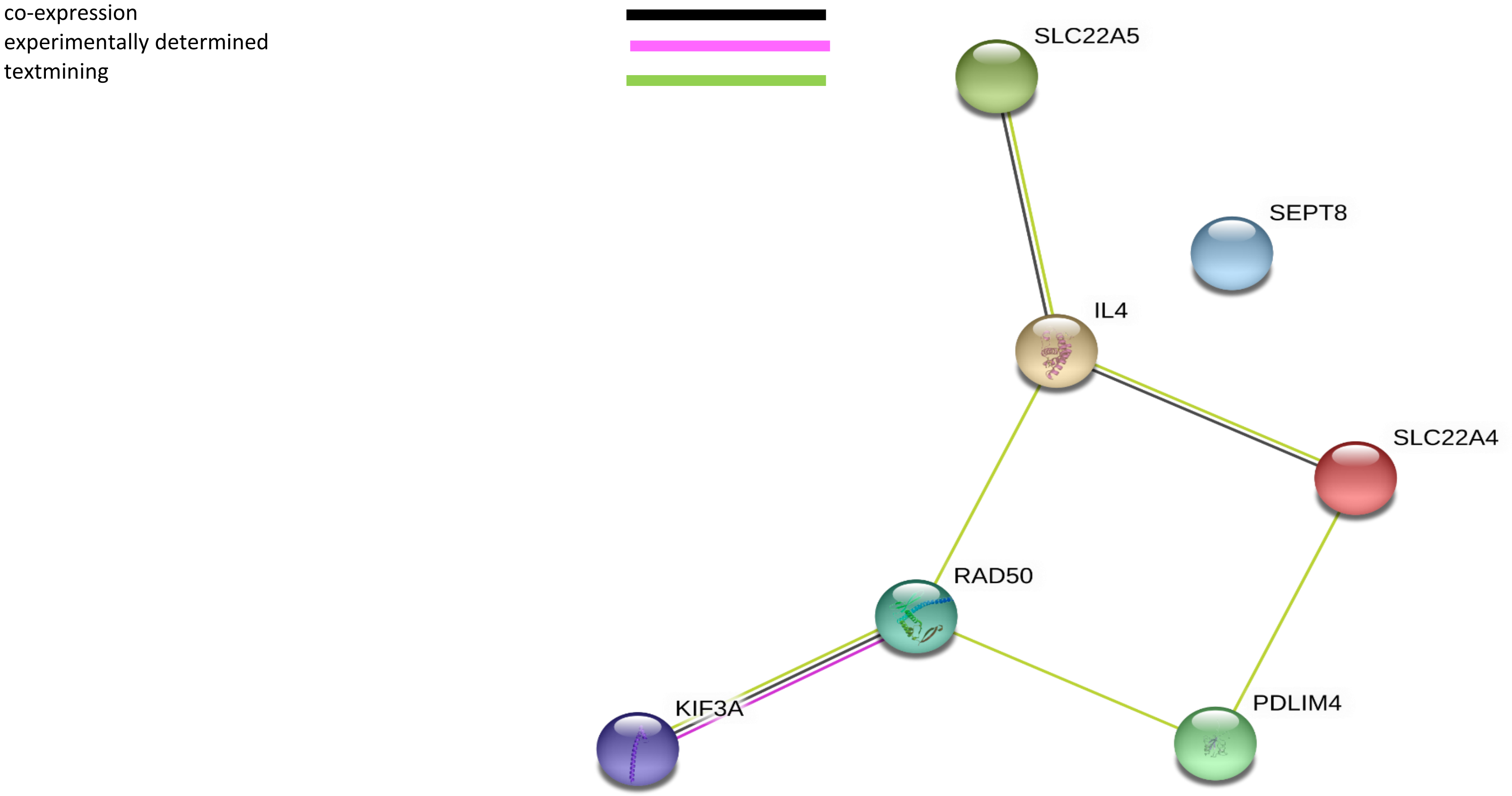

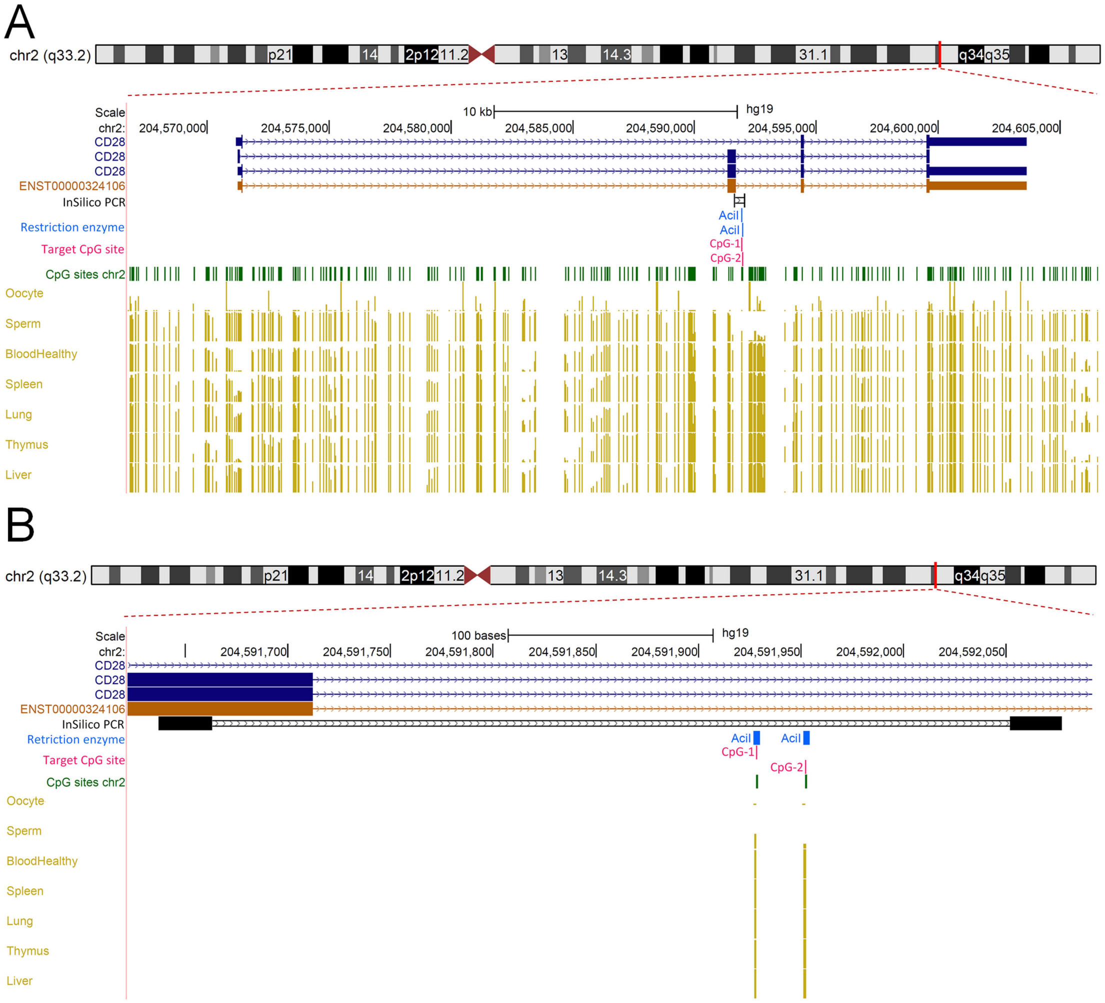

A
chr2 (q33.2)
p21
14
2p12
11.2
13
14.3
31.1
q34
q35
Scale
10 kb
hg19
chr2:
204,570,000
204,575,000
204,580,000
204,585,000
204,590,000
204,595,000
204,600,000
204,605,000
CD28
CD28
CD28
ENST00000324106
InSilico PCR
Restriction enzyme
AciI
AciI
Target CpG site
CpG-1
CpG-2
CpG sites chr2
Oocyte
Sperm
BloodHealthy
Spleen
Lung
Thymus
Liver
B
chr2 (q33.2)
p21
14
2p12
11.2
13
14.3
31.1
q34
q35
Scale
100 bases
hg19
chr2:
204,591,700
204,591,750
204,591,800
204,591,850
204,591,900
204,591,950
204,592,000
204,592,050
CD28
CD28
CD28
ENST00000324106
InSilico PCR
Retriction enzyme
AciI
AciI
Target CpG site
CpG-1
CpG-2
CpG sites chr2
Oocyte
Sperm
BloodHealthy
Spleen
Lung
Thymus
Liver

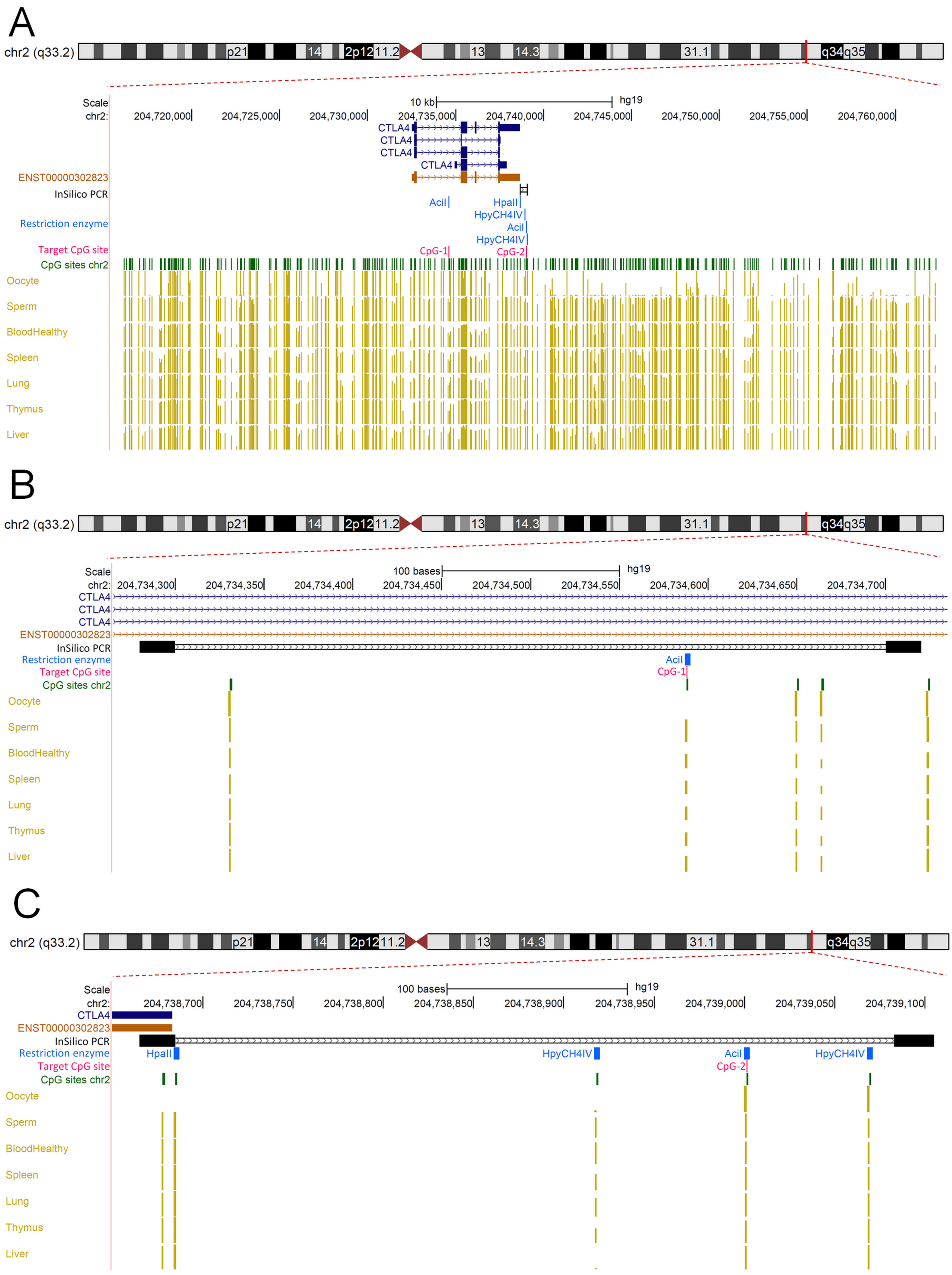
A
chr2 (q33.2)
p21
14
2p12
11.2
13
14.3
31.1
q34
q35
Scale
chr2:
10 kb
hg19
204,720,000
204,725,000
204,730,000
204,735,000
204,740,000
204,745,000
204,750,000
204,755,000
204,760,000
CTLA4
CTLA4
CTLA4
CTLA4
ENST00000302823
InSilico PCR
Restriction enzyme
AciI
HpaII
HpyCH4IV
AciI
HpyCH4IV
Target CpG site
CpG-1
CpG-2
CpG sites chr2
Oocyte
Sperm
BloodHealthy
Spleen
Lung
Thymus
Liver
B
chr2 (q33.2)
Scale
chr2:
100 bases
hg19
204,734,300
204,734,350
204,734,400
204,734,450
204,734,500
204,734,550
204,734,600
204,734,650
204,734,700
CTLA4
CTLA4
CTLA4
ENST00000302823
InSilico PCR
Restriction enzyme
AciI
Target CpG site
CpG-1
CpG sites chr2
Oocyte
Sperm
BloodHealthy
Spleen
Lung
Thymus
Liver
C
chr2 (q33.2)
Scale
chr2:
100 bases
hg19
204,738,700
204,738,750
204,738,800
204,738,850
204,738,900
204,738,950
204,739,000
204,739,050
204,739,100
CTLA4
ENST00000302823
InSilico PCR
Restriction enzyme
HpaII
HpyCH4IV
AciI
HpyCH4IV
Target CpG site
CpG-2
CpG sites chr2
Oocyte
Sperm
BloodHealthy
Spleen
Lung
Thymus
Liver

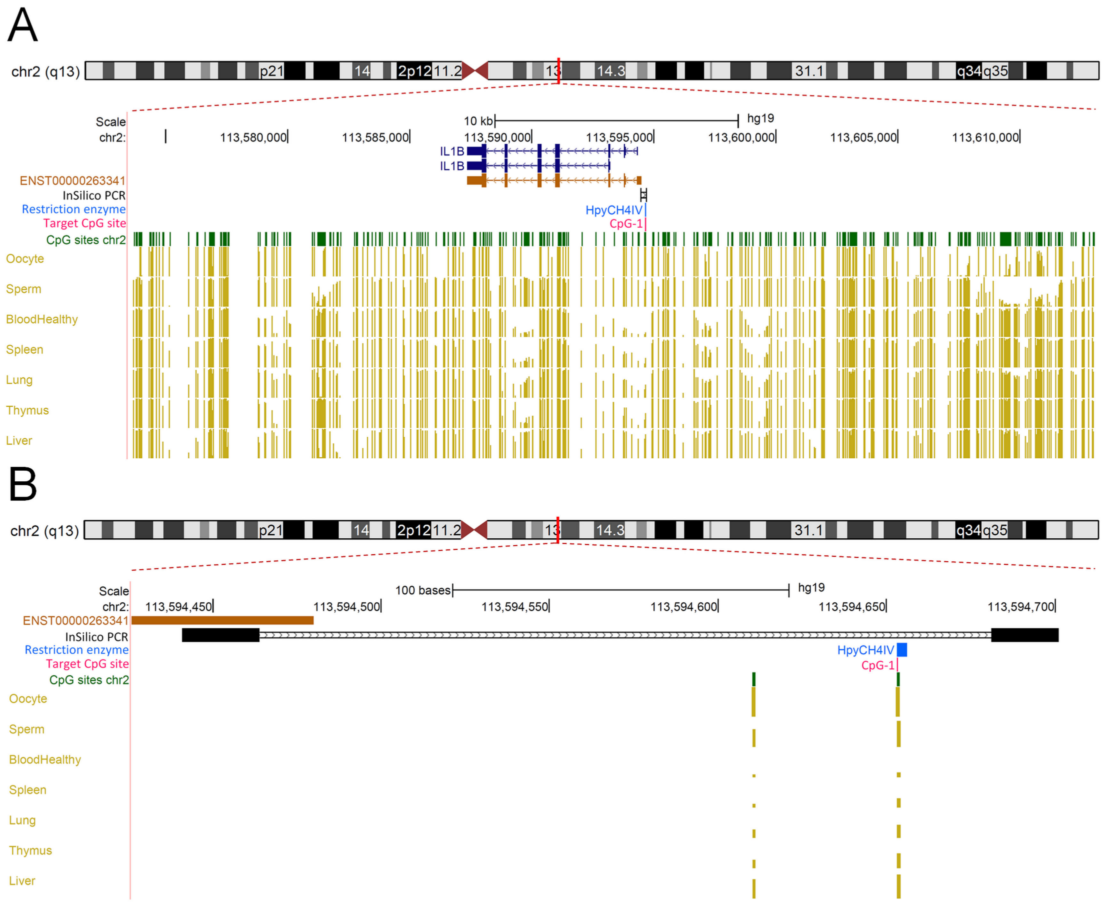

A
chr2 (q13)
p21
14
2p12
11.2
13
14.3
31.1
q34
q35
Scale
10 kb
hg19
chr2:
113,580,000
113,585,000
113,590,000
113,595,000
113,600,000
113,605,000
113,610,000
IL1B
IL1B
ENST00000263341
InSilico PCR
Restriction enzyme
HpyCH4IV
Target CpG site
CpG-1
CpG sites chr2
Oocyte
Sperm
BloodHealthy
Spleen
Lung
Thymus
Liver
B
chr2 (q13)
p21
14
2p12
11.2
13
14.3
31.1
q34
q35
Scale
100 bases
hg19
chr2:
113,594,450
113,594,500
113,594,550
113,594,600
113,594,650
113,594,700
ENST00000263341
InSilico PCR
Restriction enzyme
HpyCH4IV
Target CpG site
CpG-1
CpG sites chr2
Oocyte
Sperm
BloodHealthy
Spleen
Lung
Thymus
Liver

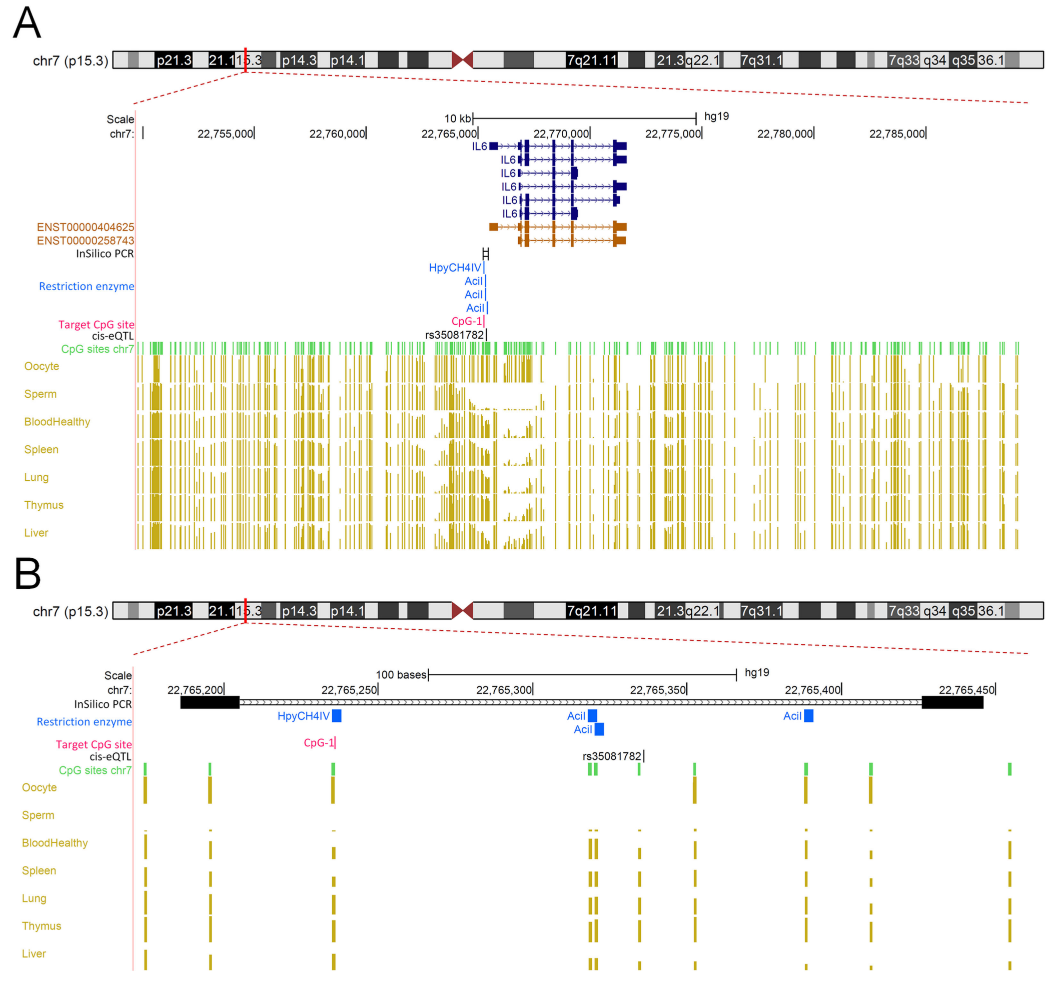

A
chr7 (p15.3)
Scale
10 kb
hg19
chr7:
22,755,000
22,760,000
22,765,000
22,770,000
22,775,000
22,780,000
22,785,000
IL6
ENST00000404625
ENST00000258743
InSilico PCR
HpyCH4IV
Restriction enzyme
AciI
Target CpG site
CpG-1
cis-eQTL
rs35081782
CpG sites chr7
Oocyte
Sperm
BloodHealthy
Spleen
Lung
Thymus
Liver
B
chr7 (p15.3)
Scale
100 bases
hg19
chr7:
22,765,200
22,765,250
22,765,300
22,765,350
22,765,400
22,765,450
InSilico PCR
Restriction enzyme
HpyCH4IV
AciI
Target CpG site
CpG-1
cis-eQTL
rs35081782
CpG sites chr7
Oocyte
Sperm
BloodHealthy
Spleen
Lung
Thymus
Liver

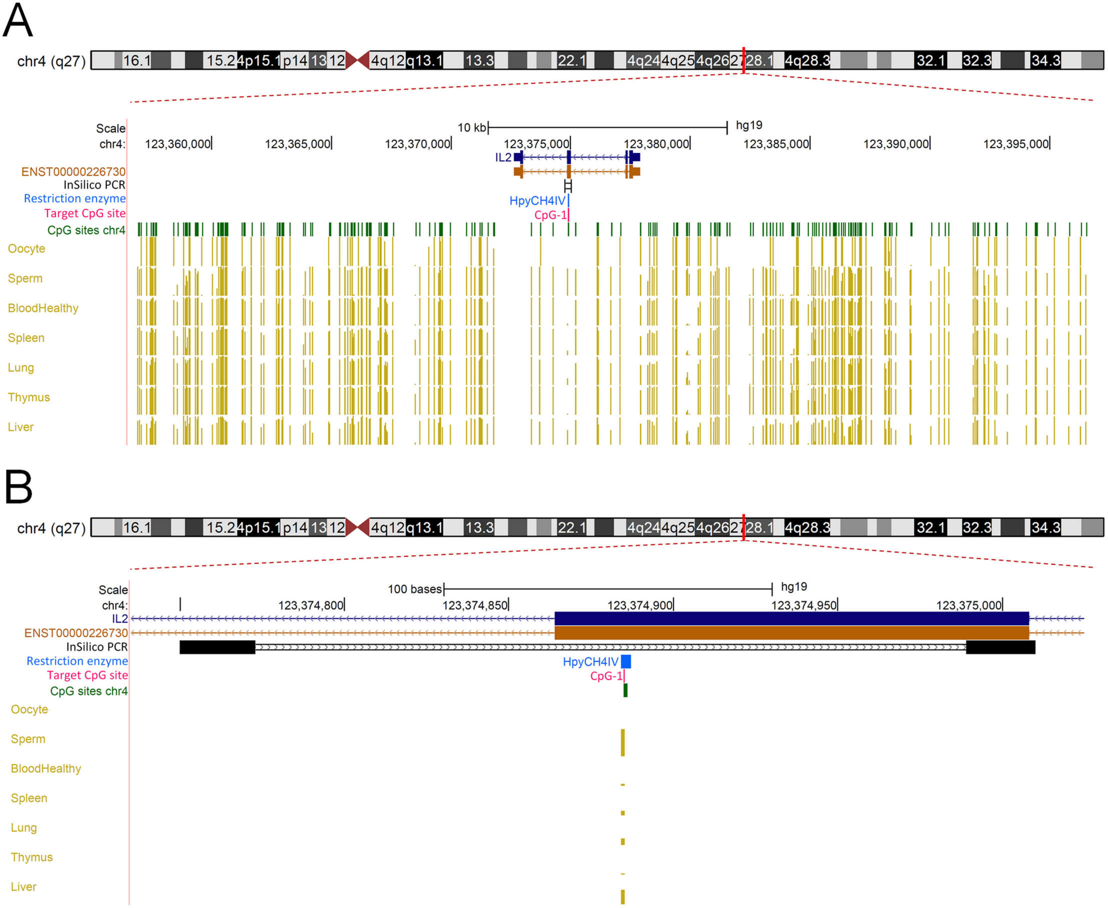

A
chr4 (q27)
Scale
10 kb
hg19
chr4:
123,360,000
123,365,000
123,370,000
123,375,000
123,380,000
123,385,000
123,390,000
123,395,000
IL2
ENST00000226730
InSilico PCR
Restriction enzyme
HpyCH4IV
Target CpG site
CpG-1
CpG sites chr4
Oocyte
Sperm
BloodHealthy
Spleen
Lung
Thymus
Liver
B
chr4 (q27)
Scale
100 bases
hg19
chr4:
123,374,800
123,374,850
123,374,900
123,374,950
123,375,000
IL2
ENST00000226730
InSilico PCR
Restriction enzyme
HpyCH4IV
Target CpG site
CpG-1
CpG sites chr4
Oocyte
Sperm
BloodHealthy
Spleen
Lung
Thymus
Liver

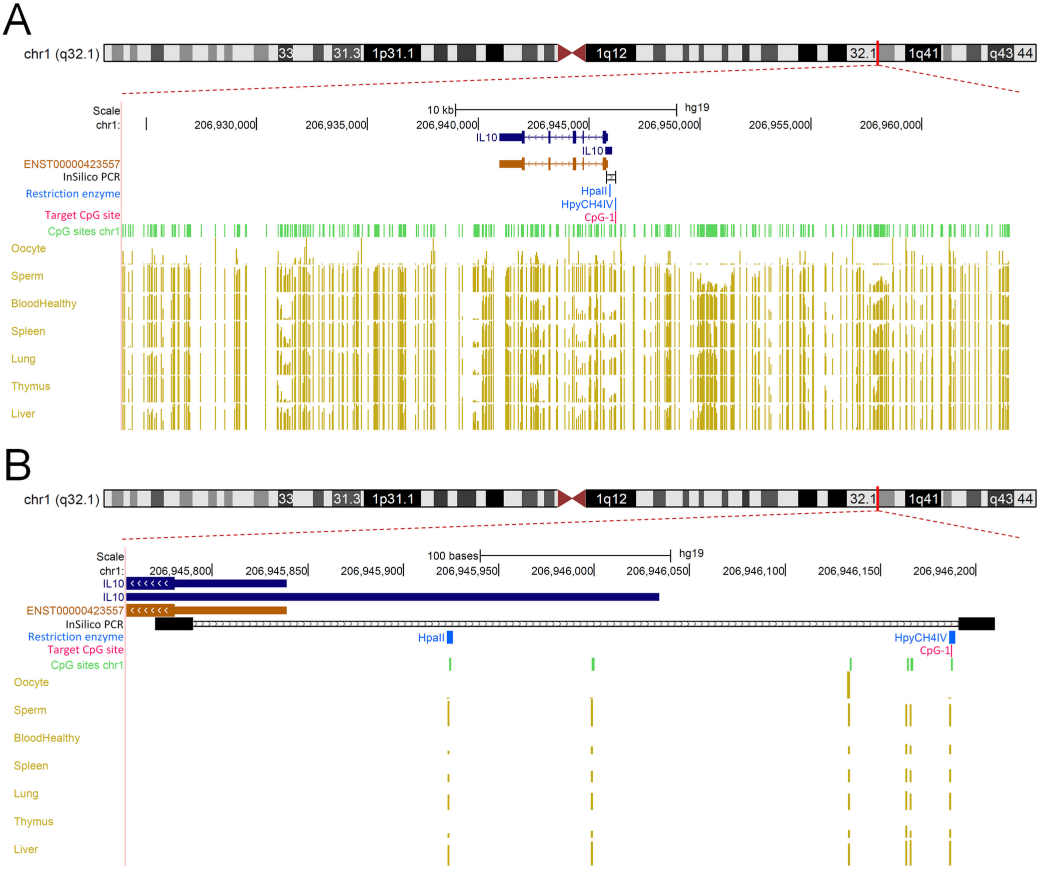

A
chr1 (q32.1)
33
31.3
1p31.1
1q12
32.1
1q41
q43
44
Scale
10 kb
hg19
chr1:
206,930,000
206,935,000
206,940,000
206,945,000
206,950,000
206,955,000
206,960,000
IL10
IL10
ENST00000423557
InSilico PCR
Restriction enzyme
HpaII
HpyCH4IV
Target CpG site
CpG-1
CpG sites chr1
Oocyte
Sperm
BloodHealthy
Spleen
Lung
Thymus
Liver
B
chr1 (q32.1)
Scale
100 bases
hg19
chr1:
206,945,800
206,945,850
206,945,900
206,945,950
206,946,000
206,946,050
206,946,100
206,946,150
206,946,200
IL10
IL10
ENST00000423557
InSilico PCR
Restriction enzyme
HpaII
HpyCH4IV
Target CpG site
CpG-1
CpG sites chr1
Oocyte
Sperm
BloodHealthy
Spleen
Lung
Thymus
Liver

A

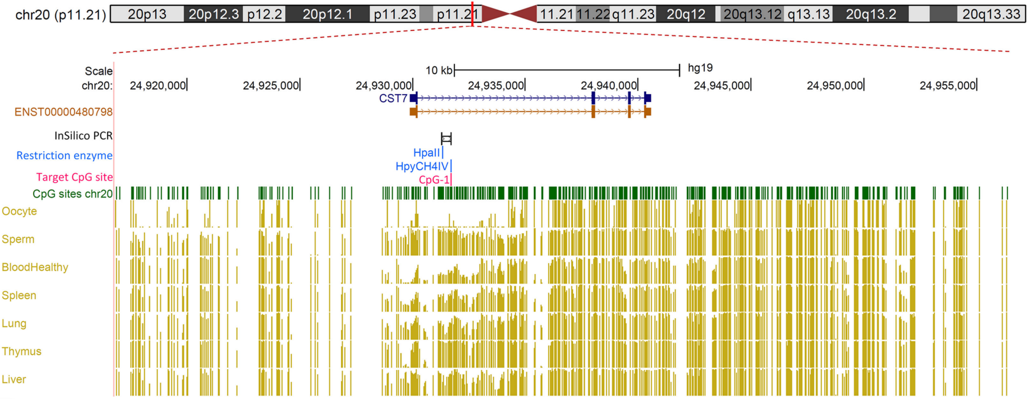
chr20 (p11.21)
20p13
20p12.3
p12.2
20p12.1
p11.23
p11.21
11.21
11.22
q11.23
20q12
20q13.12
q13.13
20q13.2
20q13.33
Scale
10 kb
hg19
chr20:
24,920,000
24,925,000
24,930,000
24,935,000
24,940,000
24,945,000
24,950,000
24,955,000
CST7
ENST00000480798
InSilico PCR
Restriction enzyme
HpaII
HpyCH4IV
Target CpG site
CpG-1
CpG sites chr20
Oocyte
Sperm
BloodHealthy
Spleen
Lung
Thymus
Liver

B

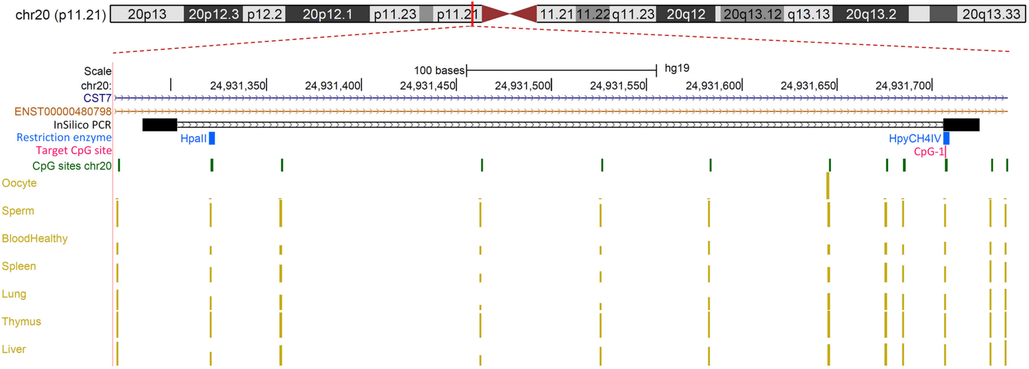
chr20 (p11.21)
20p13
20p12.3
p12.2
20p12.1
p11.23
p11.21
11.21
11.22
q11.23
20q12
20q13.12
q13.13
20q13.2
20q13.33
Scale
100 bases
hg19
chr20:
24,931,350
24,931,400
24,931,450
24,931,500
24,931,550
24,931,600
24,931,650
24,931,700
CST7
ENST00000480798
InSilico PCR
Restriction enzyme
HpaII
HpyCH4IV
Target CpG site
CpG-1
CpG sites chr20
Oocyte
Sperm
BloodHealthy
Spleen
Lung
Thymus
Liver

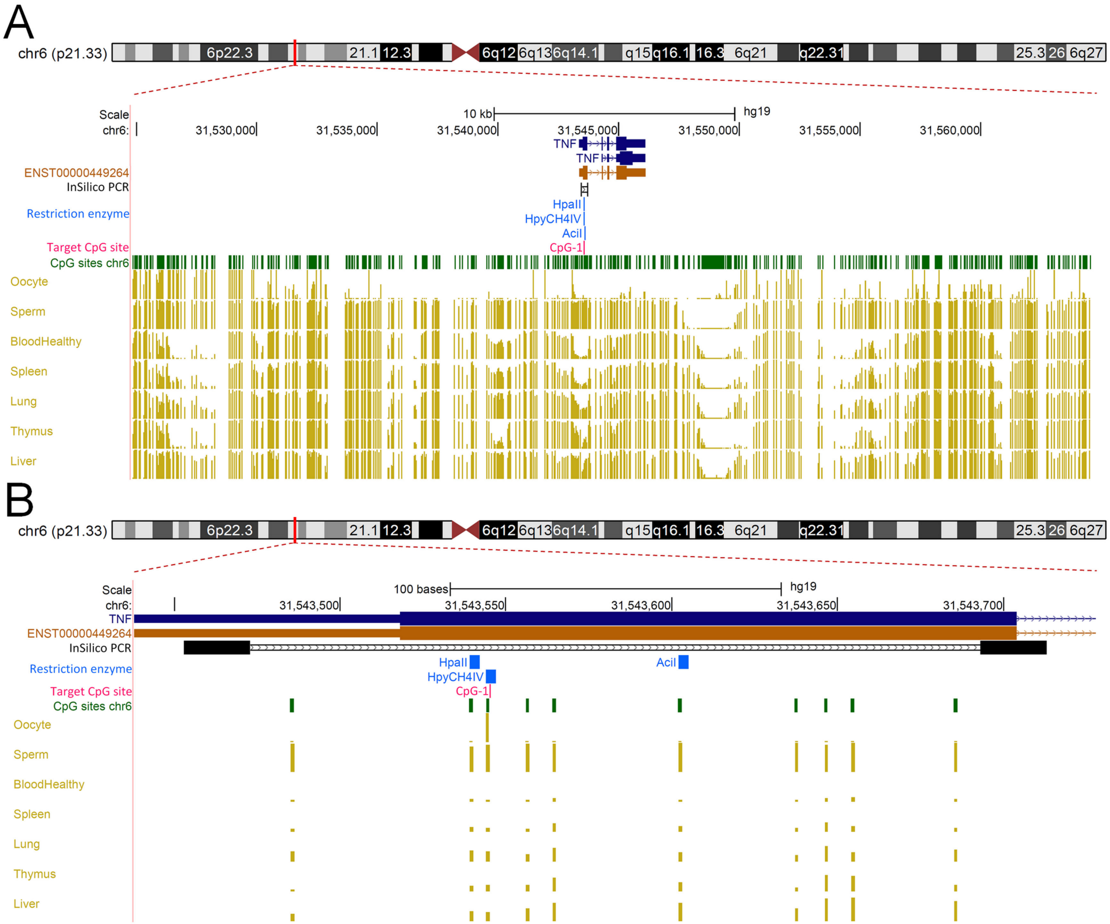
A
chr6 (p21.33)
6p22.3
21.1
12.3
6q12
6q13
6q14.1
q15
q16.1
16.3
6q21
q22.31
25.3
26
6q27
Scale
10 kb
hg19
chr6:
31,530,000
31,535,000
31,540,000
31,545,000
31,550,000
31,555,000
31,560,000
TNF
TNF
ENST00000449264
InSilico PCR
Restriction enzyme
HpaII
HpyCH4IV
AciI
Target CpG site
CpG-1
CpG sites chr6
Oocyte
Sperm
BloodHealthy
Spleen
Lung
Thymus
Liver
B
chr6 (p21.33)
Scale
100 bases
hg19
chr6:
31,543,500
31,543,550
31,543,600
31,543,650
31,543,700
TNF
ENST00000449264
InSilico PCR
Restriction enzyme
HpaII
HpyCH4IV
AciI
Target CpG site
CpG-1
CpG sites chr6
Oocyte
Sperm
BloodHealthy
Spleen
Lung
Thymus
Liver

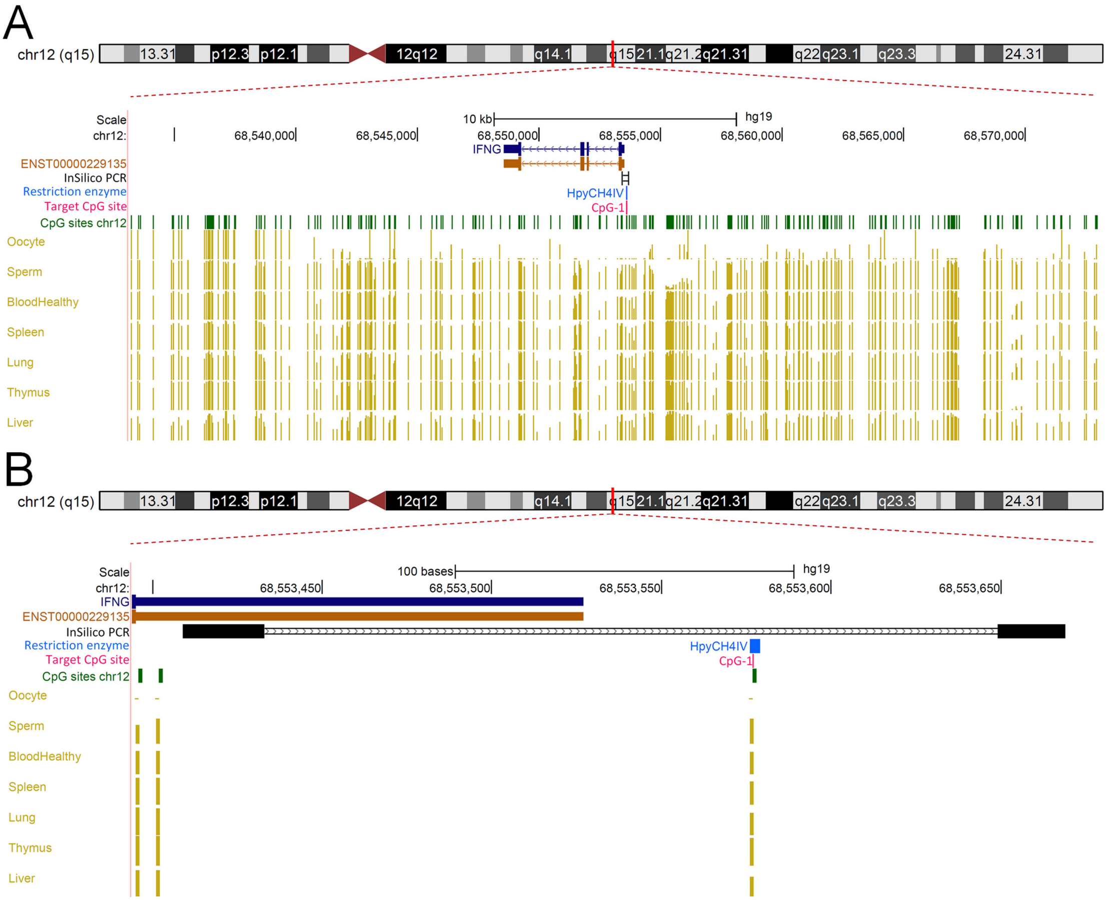
A
chr12 (q15)
13.31
p12.3
p12.1
12q12
q14.1
q15
21.1
q21.2
q21.31
q22
q23.1
q23.3
24.31
Scale
10 kb
hg19
chr12:
68,540,000
68,545,000
68,550,000
68,555,000
68,560,000
68,565,000
68,570,000
IFNG
ENST00000229135
InSilico PCR
Restriction enzyme
HpyCH4IV
Target CpG site
CpG-1
CpG sites chr12
Oocyte
Sperm
BloodHealthy
Spleen
Lung
Thymus
Liver
B
chr12 (q15)
Scale
100 bases
hg19
chr12:
68,553,450
68,553,500
68,553,550
68,553,600
68,553,650
IFNG
ENST00000229135
InSilico PCR
Restriction enzyme
HpyCH4IV
Target CpG site
CpG-1
CpG sites chr12
Oocyte
Sperm
BloodHealthy
Spleen
Lung
Thymus
Liver

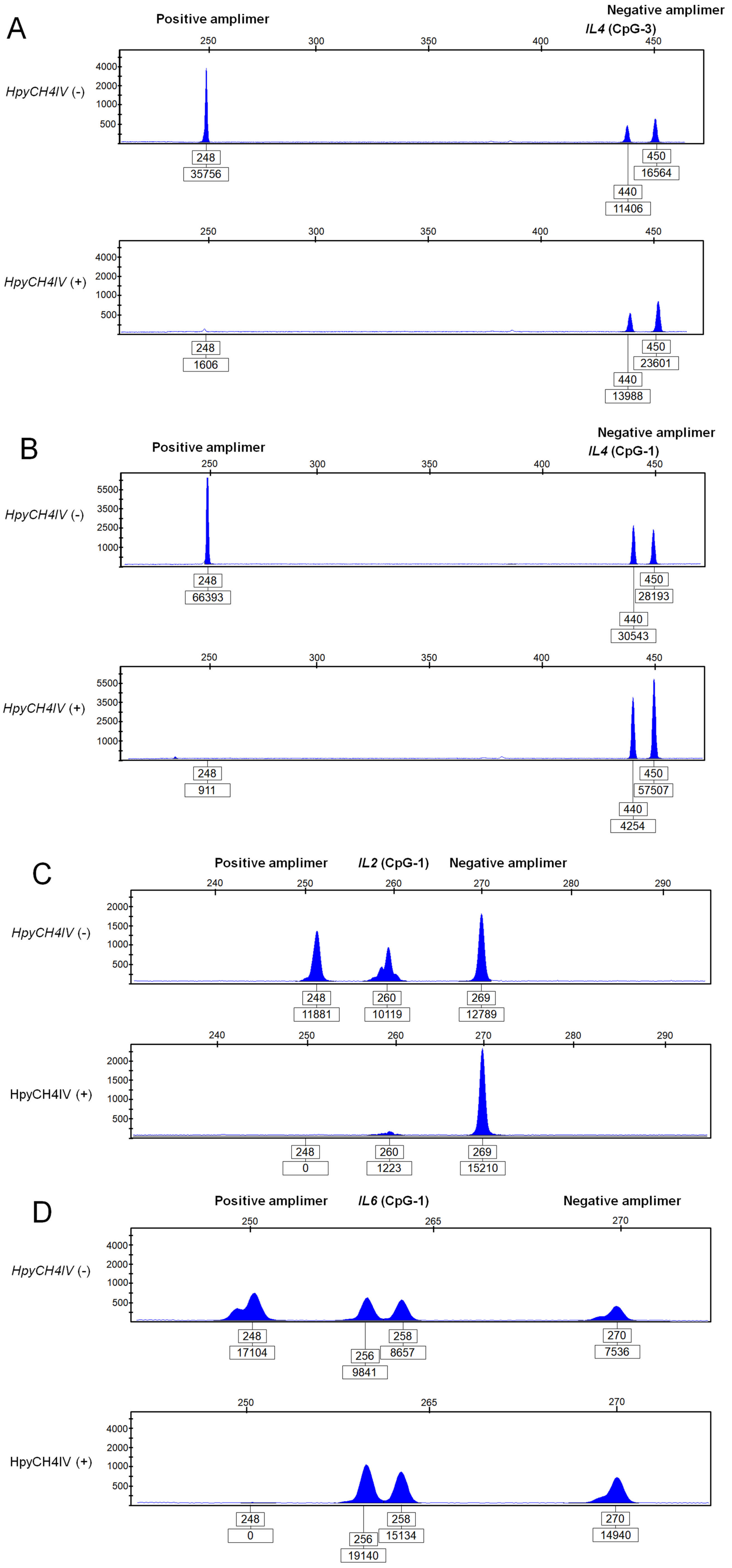

A
Positive amplimer
Negative amplimer
IL4 (CpG-3)
HpyCH4IV (-)
HpyCH4IV (+)
248
35756
440
11406
450
16564
248
1606
440
13988
450
23601
B
Positive amplimer
Negative amplimer
IL4 (CpG-1)
HpyCH4IV (-)
HpyCH4IV (+)
248
66393
440
30543
450
28193
248
911
440
4254
450
57507
C
Positive amplimer
IL2 (CpG-1)
Negative amplimer
HpyCH4IV (-)
HpyCH4IV (+)
248
11881
260
10119
269
12789
248
0
260
1223
269
15210
D
Positive amplimer
IL6 (CpG-1)
Negative amplimer
HpyCH4IV (-)
HpyCH4IV (+)
248
17104
256
9841
258
8657
270
7536
248
0
256
19140
258
15134
270
14940

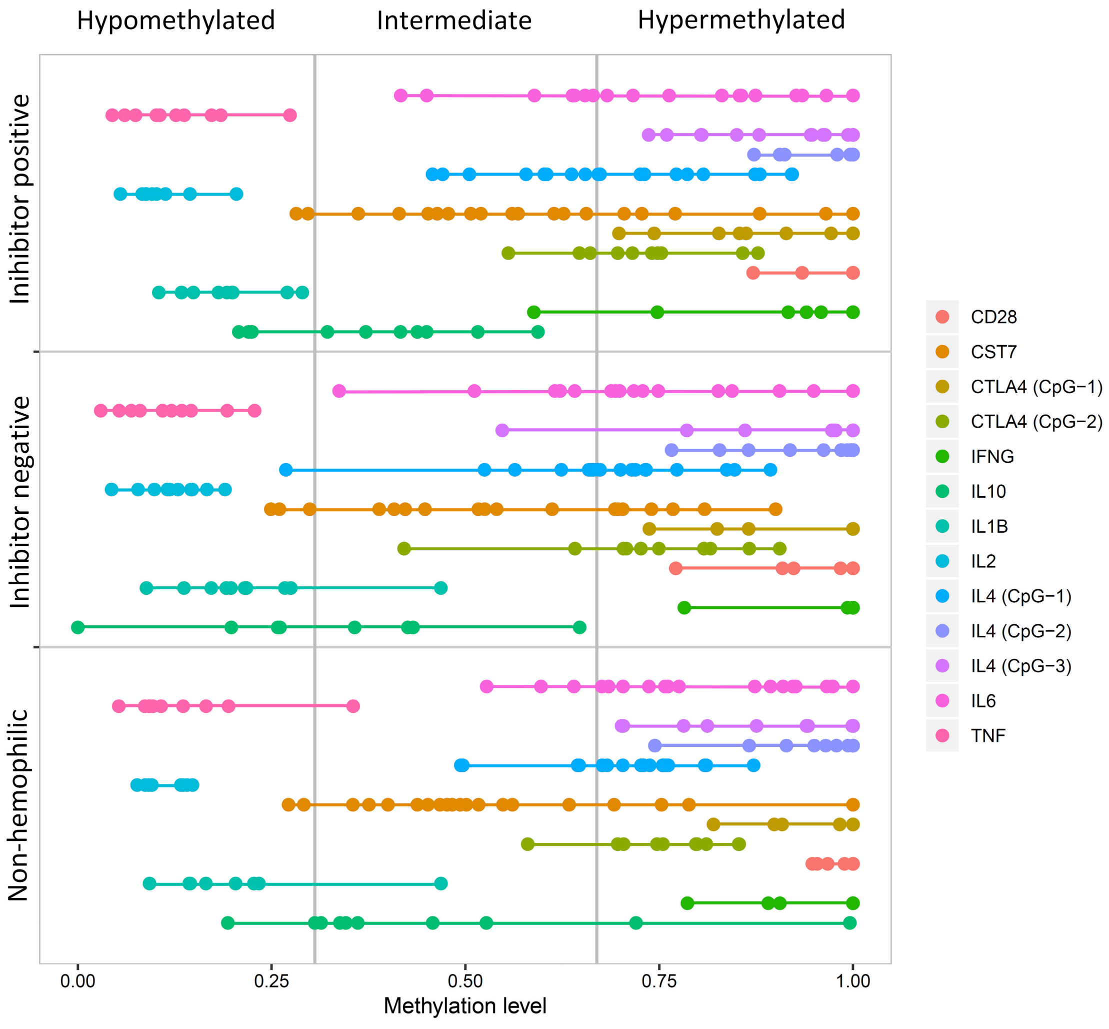

Hypomethylated
Intermediate
Hypermethylated
Inihibitor positive
Inhibitor negative
Non-hemophilic
0.00
0.25
0.50
0.75
1.00
Methylation level
CD28
CST7
CTLA4 (CpG−1)
CTLA4 (CpG−2)
IFNG
IL10
IL1B
IL2
IL4 (CpG−1)
IL4 (CpG−2)
IL4 (CpG−3)
IL6
TNF

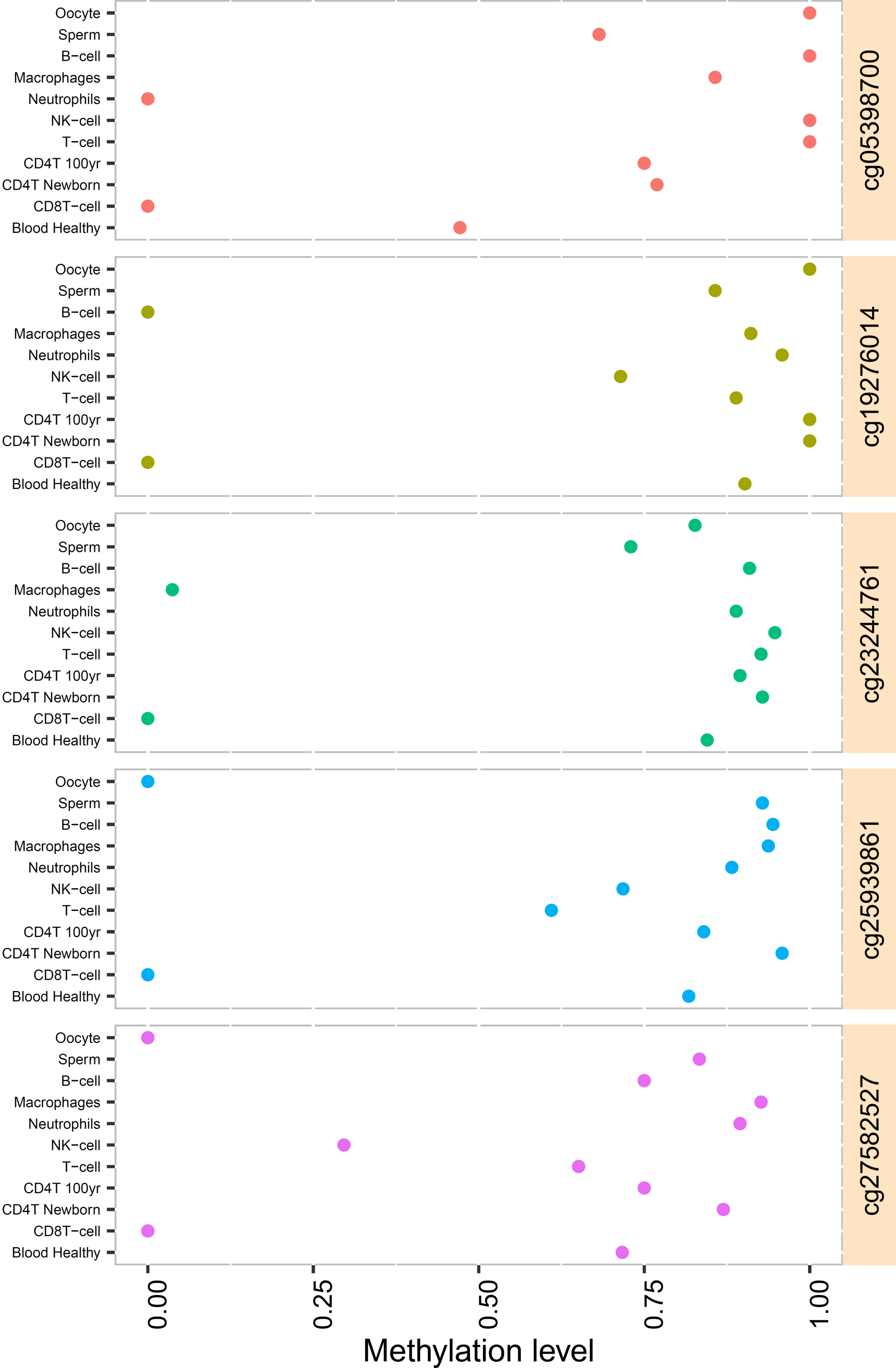
Oocyte
Sperm
B−cell
Macrophages
Neutrophils
NK−cell
T−cell
CD4T 100yr
CD4T Newborn
CD8T−cell
Blood Healthy
cg05398700
cg19276014
cg23244761
cg25939861
cg27582527
0.00
0.25
0.50
0.75
1.00
Methylation level
